# Modulating Neutrophil Extracellular Trap Formation *In Vivo* with Locoregional Precision using Differently Charged Self-Assembled Hydrogels

**DOI:** 10.1101/2024.08.16.608275

**Authors:** Tania L. Lopez-Silva, Caleb F. Anderson, Joel P. Schneider

**Affiliations:** Chemical Biology Laboratory, Center for Cancer Research, National Cancer Institute, National Institutes of Health, Frederick MD, 21702, USA

## Abstract

Neutrophil extracellular traps (NETs) are DNA networks released by neutrophils first described as a defense response against pathogens but have since been associated with numerous inflammatory diseases. The ability to induce NETs with locoregional specificity *in vivo* could facilitate studying this response in the context of disease and therapy with unprecedented control. We report the unexpected discovery that hydrogel charge predictably modulates the formation of NETs. Positively charged gels induce rapid NET release whereas negatively charged gels do not. This differential immune response to our self-assembled peptide gels enabled the development of a material platform that allows rheostat-like modulation over the degree of NET formation with anatomical and locoregional control.

## Introduction

Biomaterials that modulate specific immune responses have shown potential for therapeutic and biomedical applications,^1–3^ including vaccines,^4–6^ cancer,^7, 8^ wound healing,^9, 10^ tissue regeneration,^11, 12^ and immune tolerance.^13–15^ Commonly, materials facilitate the release of immunomodulating agents or induce an immune response by themselves based on their material properties and their interaction with immune cells upon implantation.^16^ The most well-known response to biomaterials is the foreign body reaction (FBR), where the body induces inflammation and fibrosis to encapsulate the material. However, materials that induce alternate immune responses avoiding FBR are being developed by exploring how a material’s physicochemical properties influence the immune response.^14, 17–21^ The rules governing material-immune responses are still being uncovered, and designing materials towards a targeted response remains challenging.

Neutrophils are the first responders to any injury or damage caused by the implantation of a material.^22^ The manner in which neutrophils interact with the material significantly influences the subsequent response ranging from tissue integration and repair to inflammation and FBR.^23, 24^ Thus, controlling the neutrophil response to a material offers a potential avenue in directing the subsequent immunological reaction. One such response is the formation of Neutrophil Extracellular Traps (NETs).^25, 26^ NETs are fibrous networks composed of extracellular DNA and granule enzymes such as Neutrophil Elastase (NE) and Myeloperoxidase (MPO), released upon neutrophil activation and death.^27^ Initially recognized as a defense mechanism against bacterial infection,^28^ NETs also play significant roles in inflammation,^29, 30^ autoimmune diseases,^31, 32^ cancer progression, and metastasis.^33–36^ Several factors can trigger NET formation, including bacteria, cytokines, danger-associated molecular patterns (DAMPs), lipopolysaccharides (LPS), phorbol myristate acetate (PMA), and monosodium urate crystals (MSU).^37–40^ In the context of sterile biomaterials, NET formation can be induced by the material’s properties, such as size and shape,^41^ diameter,^24^ hydrophobicity,^42^ roughness,^43^ and composition.^44^ While NET formation can be detrimental to healthy tissues and is linked to numerous pathologies, there is evidence that NETs can help resolve inflammation and protect against further tissue damage.^30^ Similarly, neutrophils and NETs have been shown to play a role in the efficacy of vaccines.^45–47^ Adjuvants like alum rely on this response to induce immunity against antigens. Therefore, developing materials that can induce NETs in a controlled manner can potentially guide the immune response and holds the promise of studying NET formation in both disease and therapy with locoregional and temporal specificity.

We are currently studying how individual design features of peptide-based gels influence the immune response. During this investigation, we designed two oppositely charged gels to define the role of electrostatic potential and made the serendipitous discovery that highly positively charged gels quickly induce the formation of NETs *in vivo*. In contrast, gels carrying negative charge do not elicit this same response. To our knowledge, the ability to induce NET formation using a gel’s electrostatic charge has not been reported and expands the list of material properties that can be exploited to induce this response. Based on this surprising observation, we developed a peptide gel platform comprising easily injected materials that can be implanted into a targeted tissue to provide site-specific anatomical control over NET formation (**Figure 1)**. Importantly, microscale locoregional control of NET release directly within a single implant is possible by modulating the distribution of charge across the material. Our gel platform can also be used to tune inflammation and dial in the degree of NET formation *in vivo* by employing composites of oppositely charged gels, **Figure 1b**. This study represents the first report of a peptide-material strategy to induce microscale NET formation *in vivo* with high locoregional control and precise tunability at a tissue and implant level.

**Figure 1.**
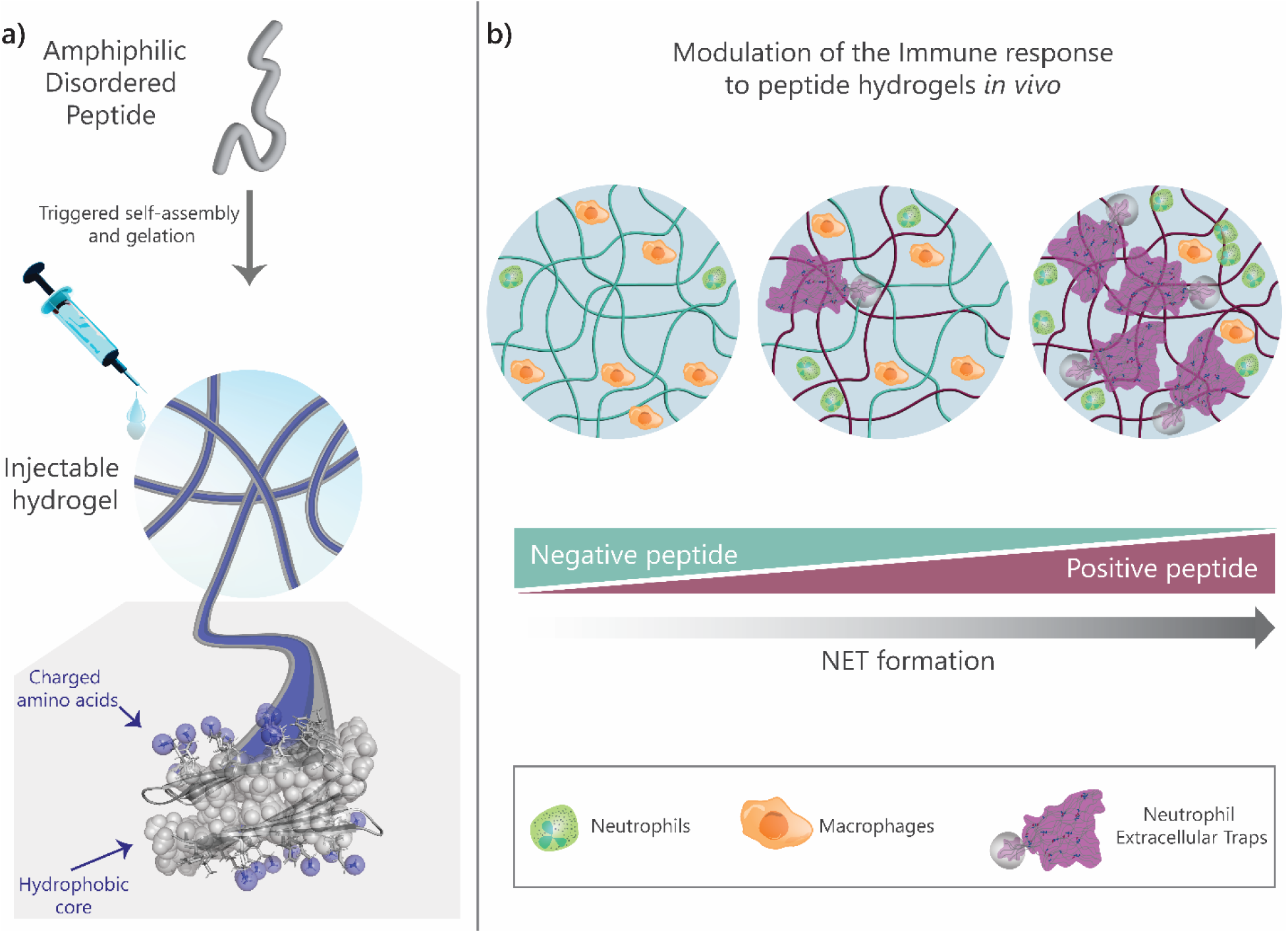
Modulating NETs formation with self-assembling peptide hydrogels. a) Hydrogels are composed of short amphiphilic peptides that self-assemble into β-sheet-rich nanofibrous networks. Implantation of peptide hydrogels by injection allows site-specific control of NET formation. b) Highly positively charged peptide hydrogels induce NET formation directly within the hydrogel implant. In contrast, negatively charged gels have initial low neutrophil infiltration and do not induce NET release. The microscale precision and degree of NET formation can be tuned by creating gradients and composites of positive and negative peptide hydrogels.

## Results and Discussion

### Designed peptides self-assemble into differently charged injectable hydrogels

Our group has developed a family of amphiphilic peptides that self-assemble into nanofibers and form hydrogels when triggered by changes in solution pH, temperature, and ionic strength. Peptides assemble into bilayered cross β-sheet fibrillar structures in which each peptide adopts a facially amphiphilic β-hairpin conformation (**Figure 1**). A solid-state NMR structure of the fibrils formed by one of our previously designed peptides showed that the hydrophobic side chains of each peptide are packed into the dry core of the fibril and the hydrophilic side chains are solvent-exposed and available for interactions with the biological milieu (**Figure 1**).^48, 49^ Herein, structure-based design was used to engineer peptides, and their respective fibrils, to direct the relative arrangement of charged residues within each peptide constituting the self-assembled fibrillar gel. This affords control over the presentation of charge and the gel’s electrostatic potential as a means to modulate the immune response. Our material design reduces biomaterial complexity as the hydrogels are made of single peptides, which carry electrostatic potential that is amplified in the final self-assembled gel network.

Figure 2a shows two β-hairpin peptides, named TLK5 and TLE5, that carry a primary charge of +5 and –5, respectively. Each peptide contains a four-residue reverse turn sequence (– V^D^PPT-) and a mixture of valine and threonine residues on their hydrophobic faces to facilitate peptide folding and hydrophobic effect-driven assembly. Importantly, each peptide also contains a block of five lysine or glutamic acid residues on their hydrophilic faces at sequence positions located near the reverse turn where the conformation of the β-hairpins is likely to be less dynamic in their assembled state. A hydrophilic non-ionic glutamine residue is also incorporated within each block. TLK5 and TLE5 peptides also contain a pair of leucine-phenylalanine residues at positions 2 and 19. These residues are known to form stabilizing pairwise interactions across β-sheets in naturally occurring proteins.^50^ Lastly, peptides are amidated at the C-termini and acetylated at the N-termini to avoid the presentation of charge at the peptide’s termini. These design features should result in peptides that undergo triggered assembly, forming gels whose fibrils display charge along their long axis (Figure 1a). All the peptides in this study were prepared by Fmoc-based solid-phase peptide synthesis, purified by RP-HPLC, and characterized by LC-MS (**Figure S1**).

**Figure 2.**
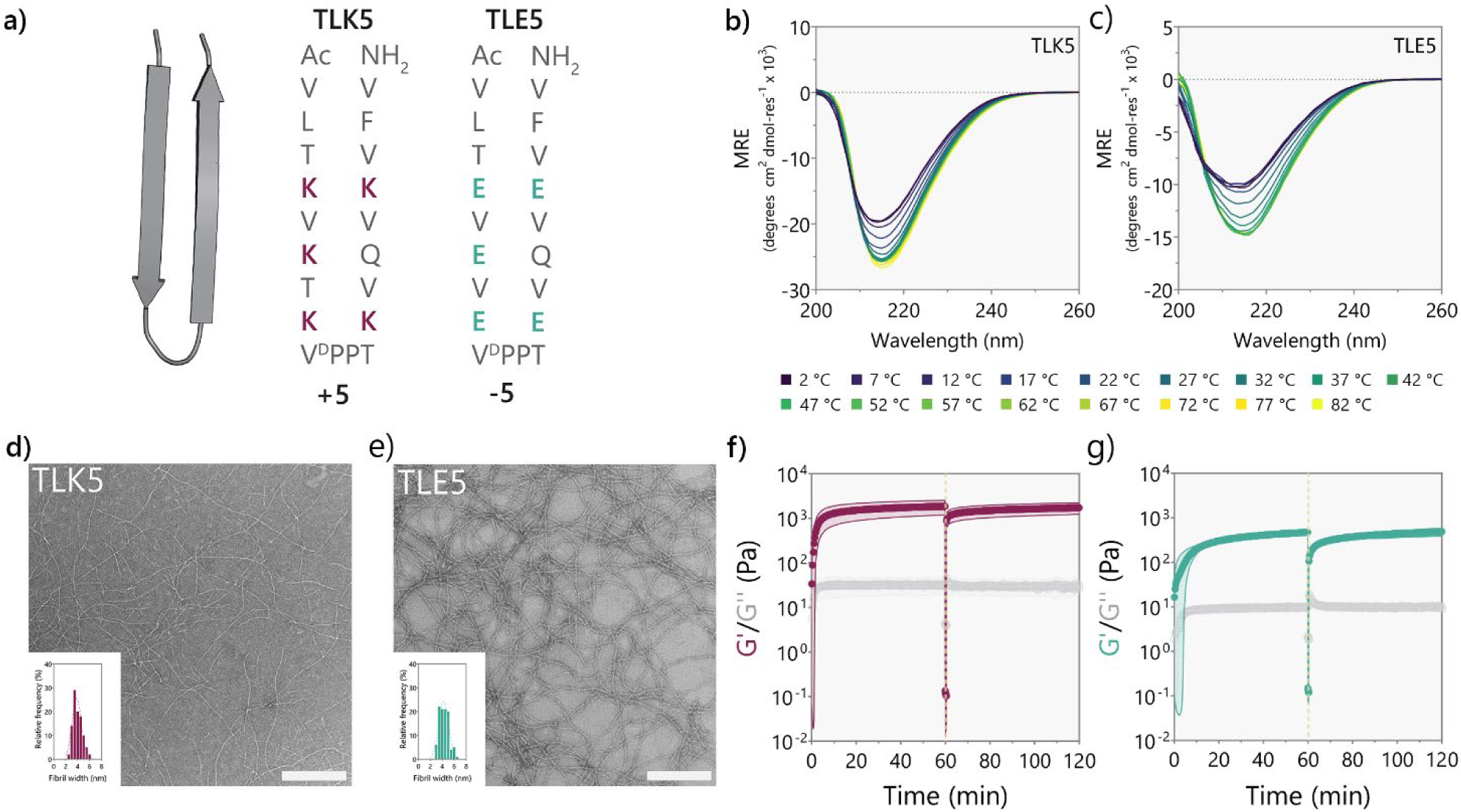
Structural and mechanical characterization of differently charged self-assembled peptide hydrogels. a) Peptide sequences for TLK5 and TLE5 shown in β-hairpin conformations. b-c) Circular dichroism spectroscopy at 10 mg/mL (1 wt.%) in 1X HBS pH 7.4 at different temperatures for b) TLK5 and c) TLE5. d-e) Negative-stain TEM images of d) TLK5 and e) TLE5 nanofibers and size distribution (inset). Scale bar represents 200 nm. f-g) Viscoelastic properties of f) TLK5 and g) TLE5 1 wt.% hydrogels in HBS at pH 7.4 over time. Hydrogels exhibit thixotropic behavior and can be injected easily, as they recover after a shear event.

The potential of TLK5 and TLE5 to assemble, fold, and form fibrillar gel networks was accessed by a combination of spectroscopy, microscopy, and rheology. Both peptides readily form 1 wt.% gels (10 mg/mL, 4.02 mM TLK5, 4.38 mM TLE5) in 1X HEPES buffered saline (HBS) at pH 7.4. Circular dichroism spectroscopy (CD) shows that both peptides adopt β-sheet structure in the gelled state, presenting spectra with distinct minima at 216 nm (Figure 2b-c).^48^ Spectra show temperature-dependent behavior consistent with assembly and β-sheet formation being driven by the hydrophobic effect.^51, 52^ Assembly is also concentration-dependent, with both peptides showing a low propensity towards β-sheet formation at lower peptide concentrations (µM range, **Figure S2-3**). TLK5 and TLE5 form long and flexible nanofibers with similar morphology as observed by TEM (Figure 2d-e). Fibrils have a diameter of around 4 nm, which is similar to the length of a folded β-hairpin and consistent with the structural model in Figure 1. Oscillatory rheology (Figures 2f-g) shows that hydrogel formation for both peptides is rapid at 37°C. TLK5 forms a gel within seconds that further cures over about 14 min (**Figure S4a**) having a plateau storage modulus (G’) of 1860 ± 667 Pa, and a loss modulus (G’’) of 32 ± 13 Pa (Figure 2f**, S5a-b and Table 1**). TLE5 forms a more compliant gel with a G’ of 470 ± 63 Pa and a G’’ of 10 ± 1 Pa that takes slightly longer to cure (Figure 2g**, S4b, S5c-d**). Both peptides form viscoelastic materials (**Table 1**) that present thixotropic behavior (Figure 2f-g), flowing under high shear stress and recovering quickly after the cessation of shear. This property allows easy implantation by injection, facilitating anatomical control over gel placement. Overall, TLK5 and TLE5 form nanofibrous hydrogels with opposite formal charges that, as will be shown, can modulate NET formation in a charge-dependent manner *in vivo*.

**Table 1.** Viscoelastic properties of peptide hydrogels.

| Peptide/Parameter | TLK5 | TLE5 |
| --- | --- | --- |
| G' (Pa) | 1860 ± 667 | 470 ± 63 |
| G'' (Pa) | 32 ± 13 | 10 ± 0.5 |
| Tan $\delta$ | 0.02 ± 0.001 | 0.02 ± 0.01 |
| Flow point (% strain) | 52 ± 12 | 88 ± 61 |
| Recovery at 5 min (%) | 71 ± 2 | 57 ± 7 |

### Differently charged gels elicit distinct cellular infiltration and degradation profiles *in vivo*

We first investigated if, indeed, material charge could influence the immune response towards TLK5 and TLE5 gels using a murine subcutaneous injection model. This model is commonly used to evaluate material biocompatibility and immune response *in vivo* because it facilitates fast screening of different materials.^53–55^ Further, for injectable gels that are minimally invasive, it provides an uninjured tissue environment minimizing additional inflammation and tissue damage.^20^ Hydrogels were harvested at 3-, 7-, 14-, and 30-days post-injection and evaluated by histology. By day 3, the positively charged TLK5 gel was infiltrated by a large number of polymorphonuclear cells, mostly at the material’s periphery (Figure 3a-c), observed as a purple ring in the H&E-stained tissue section. The peptide gel is stained pink due to its eosinophilic character provided by the lysine residues. In the infiltrated area, cells are migrating toward the core and surrounding gel fragments. By day 7, the TLK5 implants show areas of dense cellular infiltration, but the core of the gel remains unpopulated. By day 14 and 30, the implant is fully infiltrated, and the gel is being degraded, phagocytosed, and remodeled by macrophages with a characteristic foamy phenotype. Collagen is deposited within the hydrogel implant and its loose structure is similar to native subcutaneous tissue (**Figure S6**). More than 90% of the infiltrating cells are of myeloid origin (Figure 3d, **S7**), with around 70% being neutrophils (CD45^+^CD11b^+^Ly6G^+^ cells) at day 3 and 7 (Figure 3e), but by day 14 there is an increase in the percentage of monocyte/macrophage populations (CD45^+^CD11b^+^F4/80^high/low^ cells) as shown in Figure 4h-i. Two distinct populations of monocyte/macrophages are observed in TLK5 gels (Ly6C^high^ and Ly6C^low/neg^) suggesting the infiltration of macrophages with diverse phenotype, including pro-inflammatory, pro-repair, and tissue resident macrophages.^56–59^ By day 14 there is an increase of F4/80^int/low^ Ly6C^high^ cells, which have been reported to have phagocytic activity (Figure 3i). ^60, 61^

**Figure 3.**
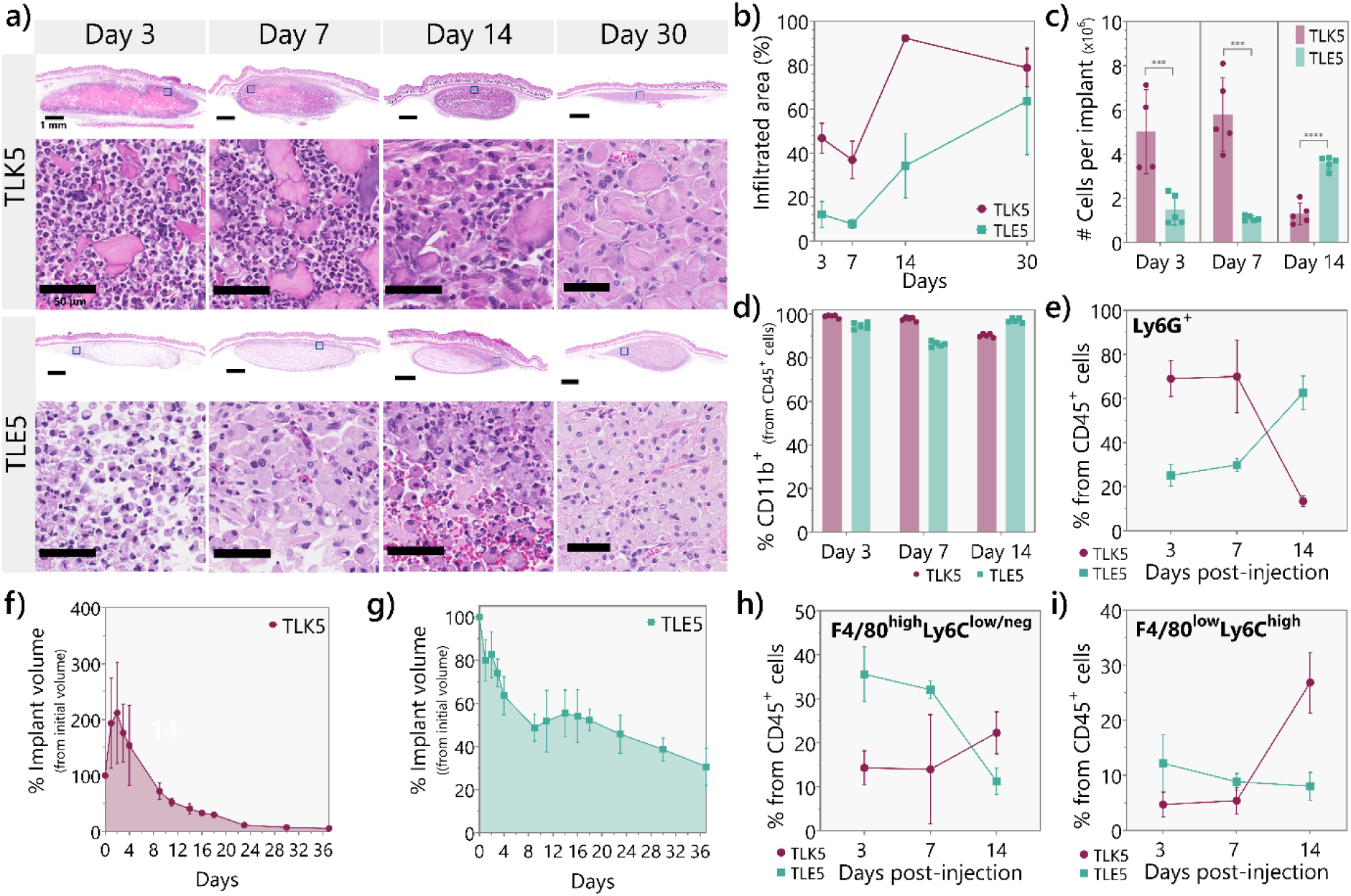
Immune response and degradation profile of TLK5 and TLE5 hydrogels in a subcutaneous injection model. a) H&E-stained tissue sections of implants at 3-, 7-, 14-, and 30-days post-injection demonstrating differential cellular infiltration patterns and evolution of the immune response. b) Percentage of cell infiltrated area over time determined by histology, n = 3 mice. Statistical differences at day 3: TLE5 vs TLK5 ** p-value < 0.003, at day 7: TLE5 vs TLK5 ** p-value 0.0037, at day 14: TLE5 vs TLK5 *** p-value 0.0004. c) Number of cells recovered per implant after digestion in preparation for flow cytometry. d) Percentage of myeloid cells (CD45^+^CD11b^+^) in the different peptide implants. e) Percentage of CD45^+^CD11b^+^Ly6G^+^ cells (Neutrophils) from total leukocytes at different timepoints. Statistical comparison: day 3: TLK5 vs. TLE5 p-value < 0.0001, day 7 TLK5 vs. TLE5 p-value < 0.0005, day 14, TLK5 vs. TLE5 p-value <0.0001, n = 5 mice. f-g) Degradation profile determined by ultrasound imaging for f) TLK5 and g) TLE5, n = 4 mice. h) Percentage of CD45^+^CD11b^+^Ly6G^neg^F4/80^high^Ly6C^low/neg^ cells from total leukocytes at different timepoints. Statistical comparison: day 3, TLK5 vs. TLE5 p-value < 0.0001, day 7, TLK5 vs. TLE5 p-value 0.0057, day 14 TLK5 vs. TLE5 p-value <0.0080, n = 5 mice. i) Percentage of CD45^+^CD11b^+^Ly6G^neg^F4/80^low^Ly6C^high^ cells (monocytes/macrophages) from total CD45^+^ leukocytes for each peptide hydrogel at 3, 7, and 14 days-post injection. Statistical comparison: day 3: TLK5 vs TLE5 p-value 0.0044, day 14: TLE5 vs TLK5 p-value < 0.0001, n = 5 mice. Error bars represent standard deviation

**Figure 4.**
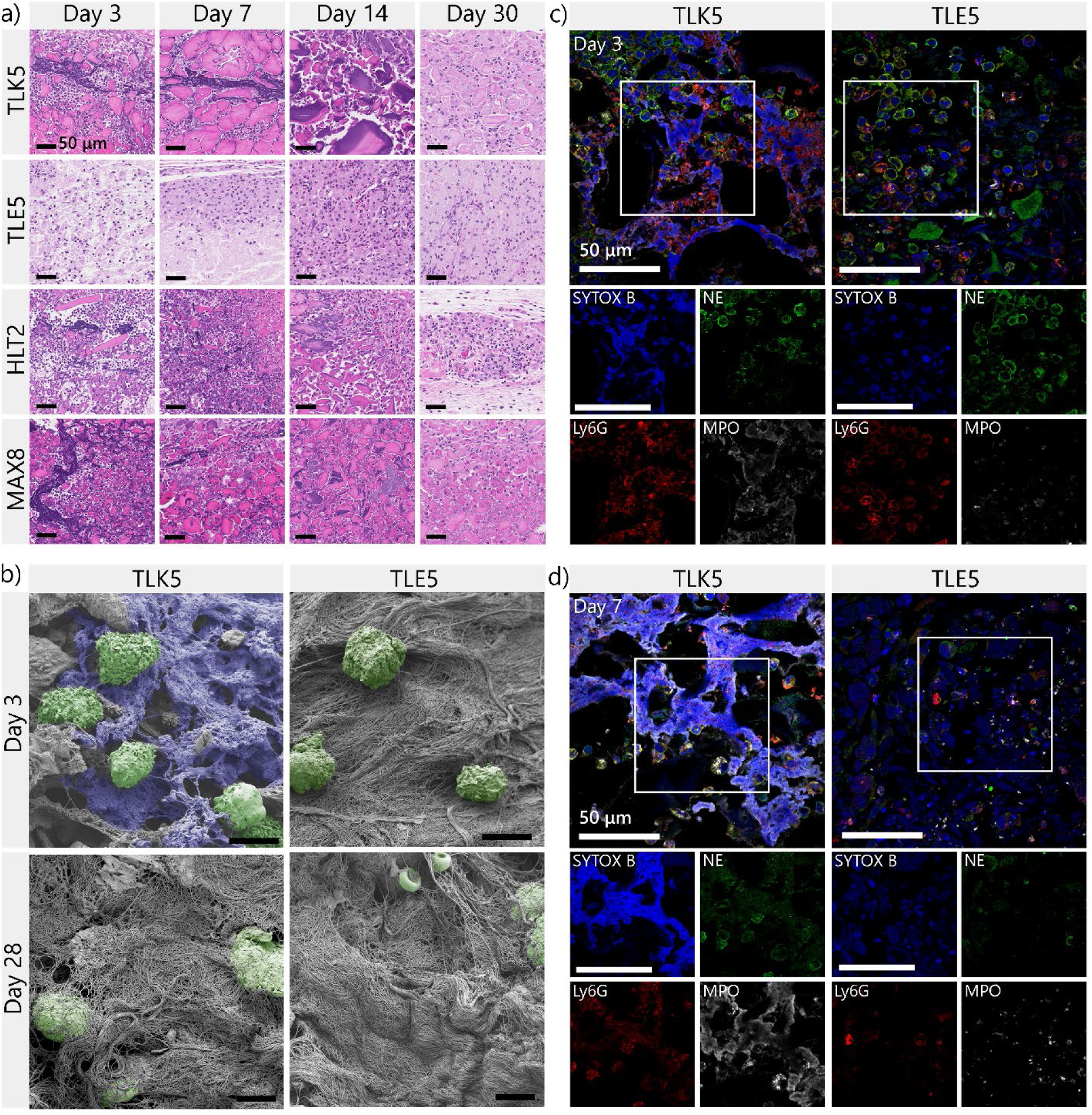
NETs formed within the implants of positively charged gels. a) H&E of implant slides at different times. The positive peptide gels have fibrous-like purple structures, indicating extracellular DNA within the implants. These DNA networks are not observed in TLE5. b) SEM images of implants to observe the NETs and implant morphology 3– and 28-days post-injection. Cells in the field of view are colored green, amorphous network structures that do not correspond to collagen or typical ECM/gel morphology are colored blue. Scale bar is 6 µm. c-d) Representative immunofluorescence images stained for NETs markers NE (green), MPO (gray), DNA (blue), and Ly6G (red) at day b) 3 and c) 7 post injection. Scale bar 50 µm.

In contrast, the negatively charged TLE5 gel had significantly fewer infiltrating cells that are located primarily at the periphery (Figure 3a-c) during the first week after implantation. The majority of these cells are macrophages with characteristic large nuclei and cytoplasm, and a F4/80 positive phenotype (Figure 3h-i). Only a few polymorphonuclear cells are observed (Figure 3e). At later time points (days 14 and 30), the TLE5 implant presents more cellular infiltration, with dense areas at the periphery of the implant surrounding a significantly less infiltrated core. Collagen is being deposited in the highly infiltrated areas (**Figure S6**). Interestingly, two weeks after injection the implant contains a higher percentage of neutrophils (Figure 3e). This is unexpected, as neutrophils are known to be early responders that attract monocytes and macrophages. However, for TLE5 gel we see that although the surface contains more macrophage-like cells (Figure 3a**, S9**), at later time points the core of the material is filled with red blood cells, neutrophils, and other immune cells. The high neutrophil content at day 14 post-injection could be due to accumulation of neutrophils from the blood present in the core of the TLE5 implants.

The presence of distinct cell types over time suggests that the degradation rate of the oppositely charged gels should differ. As such, we measured the differences in the degradation times for the TLK5 and TLE5 gels using ultrasound imaging (**Figure S10)**. This technique allowed us to follow the implant volume over the course of 38 days in the same animal with a single injection. TLK5 implants showed an increase in volume in the early days post-injection, with a peak at day 2 of around 200% from the initial volume (Figure 3f). This increase in volume is due to acute inflammation caused by the peptide hydrogel, characterized by high cellular infiltration, as seen by histology, and irritation with blood vessels surrounding the material, as observed by gross histology (**Figure S11)**. After the initial inflammation, the hydrogel volume decreased over time, with 7.3 ± 3.4 % of the initial volume remaining after a month. At 37 days post-implantation, almost all the peptide hydrogel is fully degraded and replaced with normal subcutaneous tissue with loose collagen, nerve fascicles, blood vessels, and adipose tissue (**Figure S6**). In contrast, TLE5 implants have a steady and much slower degradation rate compared with the TLK5 gel (Figure 3g). The TLE5 gel did not present the initial increase in volume, in agreement with the histological data showing minimal inflammation (Figure 3a**, S11**). In fact, 30.6 ± 8.6 % of TLE5 implant volume remains after 37 days, emphasizing the difference in degradation profiles between the two peptide gels carrying opposite charge. Overall, the TLK5 and TLE5 gels elicit divergent immune responses with different cellular infiltrates and degradation profiles, demonstrating the impact of material charge on the host immune response.

Material stiffness is known to also affect the immune response to biomaterials.^23, 62^ There is a 4-fold difference in the G’ of TLK5 versus TLE5 gels. Control studies using two previously developed electropositive peptide gels, HLT2 and MAX8 (**Figure S12**) show that charge, as opposed to stiffness, is driving the differential immune response. Both control gels are significantly softer than the TLK5 gel yet carry similar charge. Further, the G’ value of the MAX8 gel is similar to the negatively charged TLE5 gel, providing a second control. The MAX8 peptide has a formal charge of +7 and forms a gel with a G’ of around 500 Pa,^63^ whereas HLT2 (+5) forms a more compliant gel with a G’ of 233 Pa.^64^ Both MAX8 and HLT2 gels elicit a similar inflammatory response to the TLK5 gel (**Figure S13**) and degrade within the same time-frame.^63, 65^ Further, divergent responses are observed between the MAX8 and TLE5 gels, which have similar G’ values. Thus, material charge is most likely driving the immune response to the materials studied herein.

### Neutrophils release Neutrophil Extracellular Traps (NETs) within positively charged gel implants

Close inspection of the histology data at day 3 and 7 showed the unexpected formation of distinct fibrous networks at the periphery of the positively charged gels (TLK5, HLT2, and MAX8, Figure 4a**, S14-16**). These networks are formed around small gel fragments that are separated by small channels that may facilitate neutrophil infiltration. Hematoxylin staining showed that the web-like structures were basophilic, consistent with the presence of extracellular DNA, and the temporal formation of these networks correlated well with the early appearance of high numbers of neutrophils at the implant, suggesting that they could be NETs (Figure 3e). Correlation of histology, flow cytometry, and subsequent immunofluorescent staining of implant sections strongly support the assertion that these web-like structures are indeed NETs (Figure 4c-d, **S17-20**). At day 3, microscale networks of extracellular DNA (blue) co-localized with the granulocytic enzymes, myeloperoxidase (MPO, white), and to a lesser extent neutrophil elastase (NE, green). These enzymes are present in implant areas having high neutrophil density and neutrophil debris (Ly6G, red). These hallmarks of NET formation are even more pronounced at day 7, especially in the TLK5 gel (Figure 4d**, S20**). In stark contrast, no NETs were observed in the negatively charged TLE5 gel at any time point, indicating that material charge is likely the differentiating factor (Figure 4a**, S9**). TLE5 implants only display normal intracellular DNA staining within the nuclei of infiltrating cells, with a subset of these cells staining positive for intracellular NE, MPO, and membrane-bound Ly6G (Figure 4c-d, **S17-20**).

The differential immune response to the oppositely charged gels (TLK5 vs TLE5) was further examined by scanning electron microscopy (SEM). Implants were analyzed at their periphery where NETs were prominently observed by histology. At day 3, web-like networks are clearly observed in the TLK5 implant (Figure 4b**, S21**). These structures are absent in native gels that have never been implanted (**Figure S21b**). The peptide gel itself appears as a featureless grey mat by SEM in both native and implanted materials. For TLK5 implants, we also observe small channels surrounding gel fragments filled with amorphous web-like structures consistent with NET formation (**Figure S21a**). SEM also highlights the presence of infiltrating cells proximal to the observed filamentous networks. In contrast, these amorphous fibrous structures were absent in the TLE5 implant, which presented fewer cells that were surrounded by more structured collagen bundles. At day 28, amorphous networks are no longer observed in the TLK5 implant, which now contains collagen and cells. These later observations coupled with histology data suggest the resolution of NETs en route to gel degradation and tissue remodeling. Taken together, histology, flow cytometry, and SEM data suggest that the positively charged gels induce the recruitment and swarming of neutrophils upon material implantation, which interact with the material and release NETs. This early time event most likely passivates material charge and defines later immune events leading to the resolution of inflammation.

Next, we analyzed the biomolecular milieu within the implants to understand how material charge influences the production of cytokines and enzymes involved in neutrophil recruitment, activation, NET release, and the resolution of the response. In this experiment, implants were harvested over the time course of the immune response, and proteins were obtained from the implant lysates. Relevant signaling molecules required for neutrophil migration and NET formation are present at high levels in the positively charged gels early after implantation. CXCL1, a potent neutrophil chemoattractant secreted by tissue resident macrophages,^66–68^ is significantly higher in all three positive gels in comparison with the TLE5 gel at day 1 but rapidly decreases by day 3 (Figure 5a). This CXCL1 burst facilitates rapid recruitment of neutrophils immediately after gel injection, which then infiltrate the periphery of the implant, as shown by histology. A similar trend is observed for the monocyte chemoattractant CCL2 (MCP-1), the pro-inflammatory cytokine IL-6, and the Granulocyte Colony-Stimulating Factor (G-CSF), with elevated levels at day 1, followed by a decrease as the acute inflammatory response evolves (Figure 5b-d). Importantly, G-CSF has been shown to activate and prime neutrophils for NET formation, suggesting that neutrophils are activated to release NETs as early as day 1.^69, 70^ Further, increased levels of NE and MPO, granule enzymes that regulate the formation of NETs are observed at day 1 (Figure 5e-f).^28, 71, 72^ These enzymes are commonly bound to the DNA comprising NETs, which we observed by histology from day 3 through day 7. In the case of MPO, initial concentrations among groups show less pronounced differences (Figure 5f). However, TLK5 and HLT2 gels exhibit higher MPO levers at day 1 post-injection. By days 3 and 7, MPO levels are similar across all materials, which may be attributed to the presence of macrophages and other myeloid cells that produce MPO in all the gels.^73, 74^

**Figure 5.**
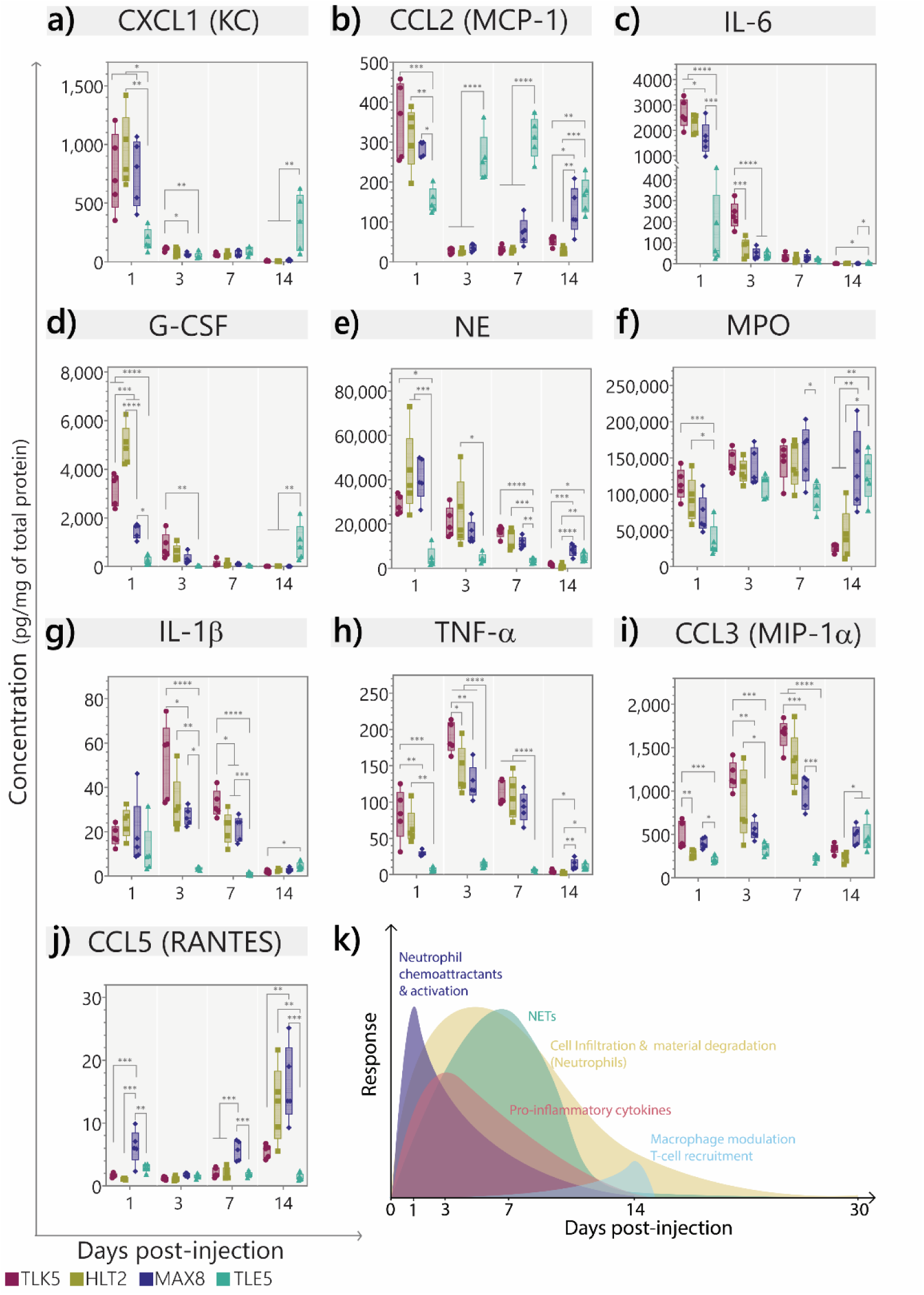
Evolution of signaling molecules involved in neutrophil recruitment and NET formation. a-j) Concentration of enzymes, cytokines, and chemokines implicated in neutrophils recruitment, activation, and NET formation at 1, 3, 7, and 14-days post-injection. Data are represented as range with minimum and maximum values and the mean is represented as a line. n = 5. Statistical comparisons are shown in the supporting information. k) Representation of the immune response elicited by TLK5.

Levels of the pro-inflammatory cytokines IL-1β and TNF-α are moderate in the positively charged gels at day 1 but increase and peak at day 3 (Figure 5g-h). We also observe high concentrations of the macrophage inflammatory protein-1α (MIP-1α) that peaks at day 7 (Figure 5i). Neutrophils produce this chemokine as a chemotactic for monocytes and neutrophils, and it is relevant for the progression of the inflammatory response elicited by the positive gels.^75^ In fact, we see an increase in the monocyte/macrophage percentage at day 14 for these materials, possibly due to the release of MIP-1α by neutrophils. All of these cytokine levels decrease with the concomitant increase of CCL5 at day 14. CCL5 is involved in lymphocytes and macrophage reprogramming to the resolution phase of inflammation (Figure 5j).^76^ Taken together, this suggest that the positively charged materials induce an acute inflammatory response that resolves over time. In contrast, the negatively charged TLE5 gel exhibited lower levels of all these biomarkers at day 1. Further, cytokines associated with acute inflammation remain low over the course of the study. Levels of MCP-1 progressively increase and peak around days 3-7, which correlates with the presence of macrophages as seen by flow cytometry. Thus, the negatively charged gel does not induce the same degree of inflammation, and importantly, the infiltrating neutrophils observed on day 14 do not produce NETs (**Figure S21**).

Figure 5k summarizes the temporal immune response to the positively charged gels correlating all the data collected thus far. These gels elicit an acute inflammatory response immediately after injection that is characterized by a fast recruitment and activation of neutrophils, the formation of NETs observed at day 3 and 7, the recruitment of monocytes and macrophages as the response progresses. Importantly, we observe the resolution of inflammation accompanied by gel degradation and elimination of NETs within a month (Figure 5k). Thus, for these positively charged gels, NET formation is not followed by chronic inflammation, tissue damage, or the FBR. The differential immune response observed as a function of material charge opens the possibility of developing a material platform capable of controlling the location and degree of NET formation.

### Locoregional control of NET formation

The contrasting immune responses triggered by the oppositely charged TLK5 and TLE5 gels allows precisely control of NET formation within a single implant. The distribution of charge can be controlled by taking advantage of the rheological properties of these gels. Separate TLK5 and TLE5 gels can be sequentially drawn into a syringe and then delivered to a common site with minimal mixing (Figure 6a). This results in the delivery of a single implant with two separate domains: one containing TLK5 enriched with positive charge and the other enriched with negative charges provided by TLE5. Once implanted, distinct immune responses are elicited by the spatially distinct TLK5 and TLE5 domains (Figure 6b). One side of the implant has high cellular infiltration by polymorphonuclear cells and contains NETs, similar to the immune response observed in implants of TLK5 alone (Figure 6b-c). Conversely, there is lower cell infiltration and absence of NETs in the TLE5-containing domain (Figure 6b-d). Thus, inflammation and NET formation can be controlled with locoregional specificity by simply engineering the delivery of two oppositely charged materials.

**Figure 6.**
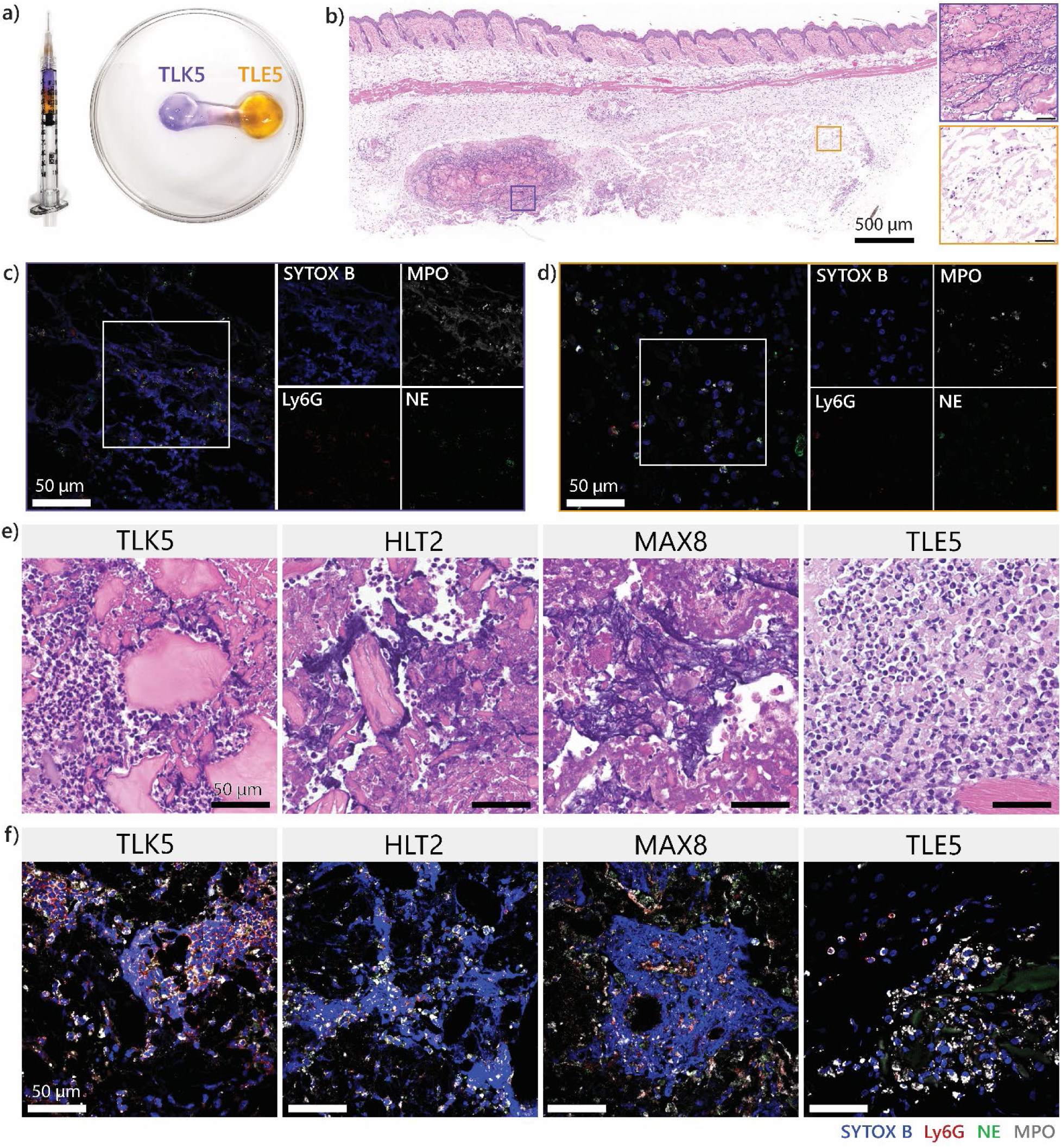
Locoregional control of NET formation. a) Charge distribution within a single implant is controlled by sequential loading and injecting TLK5 and TLE5 gels via syringe. b) H&E-stained tissue section of the resulting implant containing two domains of oppositely charged gels 3 days post-injection. The inset outlined in purple shows a region of the TLK5 domain. The orange inset shows a region enriched with the TLE5 gel. c) Immunofluorescent staining for DNA, Ly6G, NE, and MPO shows that NETs formed in the TLK5 domain. d) Immunofluorescent staining of the TLE5 domain. e) H&E-stained tissue sections of peptide gels injected in the gastrocnemius muscle showing the presence or absence of NETs 3 days post-injection. f) Merged immunofluorescent images of intramuscularly injected gels stained for NET markers.

Anatomical control of NET formation is possible by the facile delivery of the gels into a tissue of interest by simple injection. We performed intramuscular injections of all the gels into the mouse gastrocnemius muscle and assessed NET formation by histology at day 3 (**Figure S23**). All the materials were highly infiltrated by cells, but only the positively charged gels contained areas with NETs as confirmed by hematoxylin (Figure 6e) and immunofluorescent staining (Figure 6f). NETs are only observed within the implant and do not form in the muscle fibers, demonstrating microscale locoregional control over the response (**Figure S24**). However, the muscle around the gel implant appears to have more cellular infiltration than the control muscle without injection (**Figure S24**). Although histology suggest that the immune response elicited in the muscle and the subcutaneous space are likely not identical,^55^ neutrophil recruitment and NET release are conserved in both anatomical sites early in the immune response. It would certainly be interesting to assess if this is the case in other tissue types, an area of future work.

### Controlling the degree of NET formation by modulating material charge

Given that material charge is a key determinant of the neutrophil response, we hypothesized that the degree of inflammation and NET formation could be tuned by adjusting the overall charge displayed within the gel network. To test this hypothesis, we prepared composite materials where charge was systematically varied by first preparing separate TLK5 and TLE5 gels and then combining them at different volume ratios (100, 75, 50, 25, and 0% TLK5 relative to TLE5) by shear-thin mixing. This results in gel composites with a range of positive charge content. Each gel composite was injected subcutaneously, and the immune response evaluated at day 3 when NETs were first observed for the TLK5 gel. Notably, histology showed that fine control over the degree of inflammation could be achieved by modulating the material charge (Figure 7a). Gross histology images show a spectrum of implant appearances that range from the characteristic irritation and yellow coloration, previously observed for the 100% TLK5 gel, to a clear implant as seen before for the 100% TLE5 gel. We also observe a rheostat-like variation on the degree of cellular infiltration by H&E staining (Figure 7b). The composite with 75% TLK5, which is predominantly positively charged, shows a similar infiltration profile to pure TLK5, where polymorphonuclear cells infiltrate into the periphery of the implant using channels that surround gel fragments. In contrast, composite with 25% TLK5 has lower cell infiltration and resembles more the pure TLE5 implants. The composite comprised of an equal mixture (50%) of TLK5 and TLE5 has a distinct appearance, with areas of high cellular infiltration mixed with areas with low cell density (Figure 7b). This suggest that the implant displays charge heterogeneously, having distinct microscale domains of positive and negative charge. This heterogeneity is probably a reflection of the mixing efficiency of the two gels. We observed similar trends with respect to NET formation. NETs are observed in the 100, 75, and 50 % TLK5 composites by immunofluorescent staining (Figure 7c). NETs are formed at the periphery of the 75% TLK5 composite, similar to the pure TLK5 gel. Interestingly, for the 50% TLK5 composite, NET formation appears to take place only in the microscale domains that contain high cell density (Figure 7b). Here DNA networks are confined into distinct regions that are surrounded by domains with fewer cells and no NETs. Lastly, no NETs were observed in the in 0% and 25% TLK5 composite gels.

**Figure 7.**
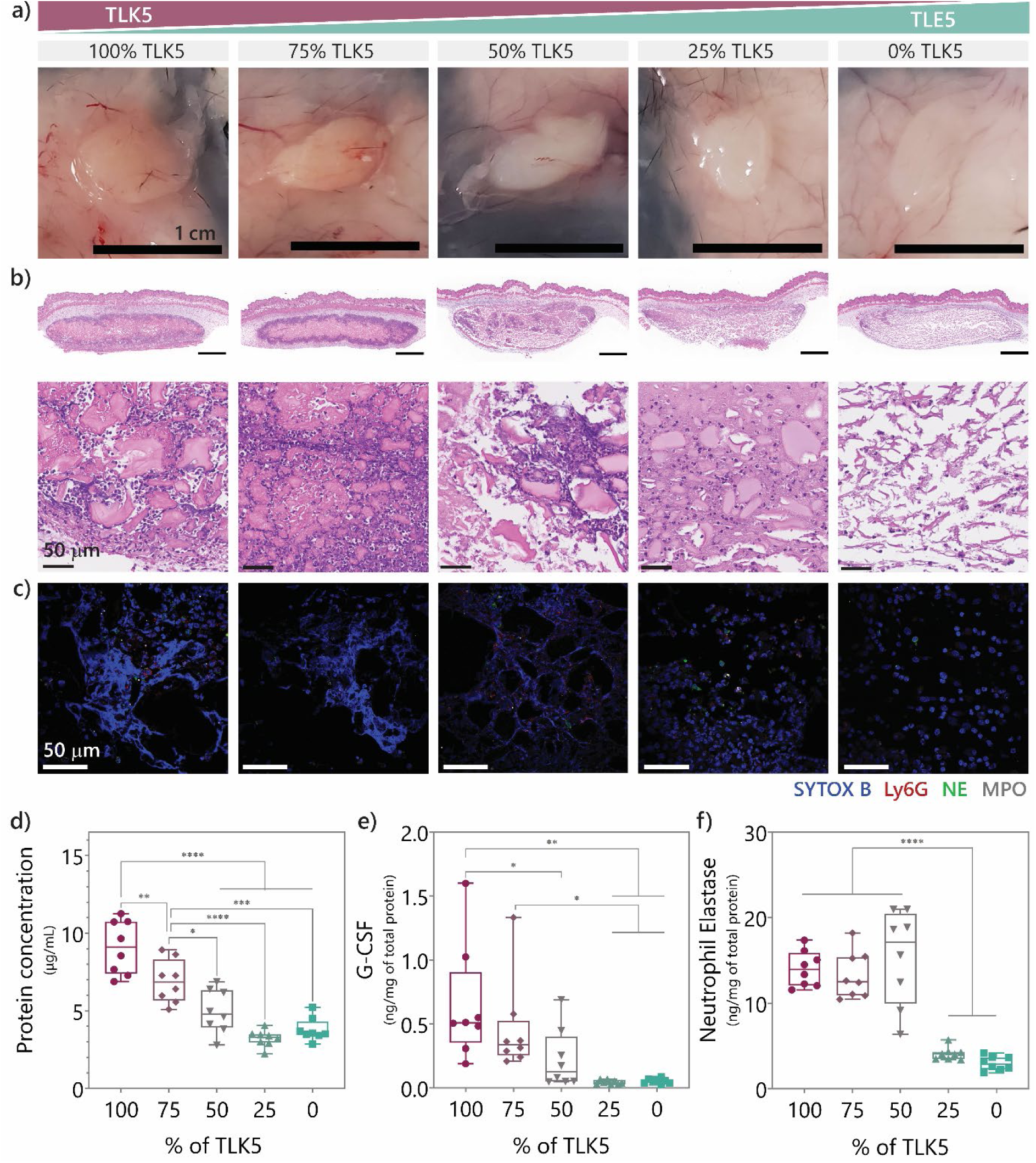
Composites gels of varying charge enable fine-tuning of the degree of inflammation and NET release. a) Gross histology of implants of TLK5-TLE5 composites ranging from 100% TLK5 to 0% TLK5 3 days post-injection. b) H&E-stained tissue sections of the different composites implanted subcutaneously. c) Immunofluorescent-stained images of the different implants for DNA, Ly6G, NE, and MPO. d) Protein concentration of lysates from the different composites. Data are represented as range with minimum and maximum values and the mean is represented as a line. n = 8. Statistical comparisons * p-value: 0.0301, ** p-value 0.0096, *** p-value: 0.0001, **** p-value <0.0001. e) Concentration of Neutrophil Elastase present in the implant lysates with different TLK5 content. Data are represented as range with minimum and maximum values and the mean is represented as a line. n = 8. Statistical comparisons **** p-value <0.0001. f) Concentration of G-CSF in the implant lysates with a range of TLK5 percentages. Data are represented as range with minimum and maximum values and the mean is represented as a line. n = 8. Statistical comparisons, 100% vs 50% * p-value:0.0383, 100% vs 25% ** p-value 0.0013, 100% vs 0% p-value 0.0015, 75% vs 25% p-value 0.0416, 75% vs 0% p-value 0.0484.

This rheostat behavior is also observed at the level of secreted protein and NET biomarkers measured from implant lysates isolated at day 3. Total protein, G-CSF and NE, which are involved in neutrophil activation and NET regulation, vary according to the amount of TLK5 gel present in each composite. Total protein levels, an indicator of cell infiltration, decrease in nearly a stepwise fashion as the amount of TLK5 in each composite is decreased (Figure 7d). With respect to neutrophil activation, composites containing higher amounts of TLK5 gel (50-100%) have higher levels of G-CSF (Figure 7e). Of note, in this experiment, G-CSF levels are measured at day 3. Higher levels most likely exist at earlier times, as was observed for the pure TLK5 gel at day 1 (Figure 5d). This suggests that priming of neutrophils to release NETs may take place early for the composite gels as well. Composites with higher TLK5 content (50-100%) also had higher levels of NE in comparison to the 0-25%TLK5 composites (Figure 7f). As expected, NETs are observed by histology in composite implants having higher concentrations of NE (Figures 7b**,c****,f**). These data demonstrate that formulating composite gels of varying positive charge afford control over the degree of inflammation and NET formation.

## Conclusion

Herein, we found that charge can be used as a design element in the construction of peptide gels that predictably control NET formation *in vivo*. The highly positively charged gel formed by the self-assembling peptide TLK5 induces an acute inflammatory response defined by the recruitment and infiltration of neutrophils that then release NETs within the implanted material. Over time, inflammation resolves, and the gel and NETs are cleared by infiltrating macrophages. In contrast, the negatively charged gel formed by peptide TLE5 elicited low cellular infiltration and no NET formation. These two materials, with divergent immune responses, serve as the foundation of a material platform that enables precise spatiotemporal control over NET formation. This platform allows anatomically specific NET formation, demonstrated here for subcutaneous and intramuscular tissues. NET formation can even be controllably induced within specific regions of a single implant via the differential presentation of opposite charge. Further, the degree of inflammation and NET formation can be dial in by employing composite gels of varying positive charge. This material-based strategy affords unprecedented control over NET release *in vivo* and offer the promise of studying NET formation in both disease and therapy with locoregional specificity.

## Associated Content

The Supporting Information is available free of charge at

Supplementary Figures S1-24, Tables S1-S3.

Materials and Methods, additional data for peptide purity and characterization, flow cytometry panel, histological data, SEM data, and statistical analyses.

## Author Information

### Corresponding author

**Joel P. Schneider** – *Chemical Biology Laboratory, Center for Cancer Research, National Cancer Institute, National Institutes of Health, Frederick, MD, 21702, United States*.

### Authors

**Tania L. Lopez-Silva** – *Chemical Biology Laboratory, Center for Cancer Research, National Cancer Institute, National Institutes of Health, Frederick, MD, 21702, United States*.

**Caleb F. Anderson** – *Chemical Biology Laboratory, Center for Cancer Research, National Cancer Institute, National Institutes of Health, Frederick, MD, 21702, United States*.

## Acknowledgments.

This work was supported by the National Cancer Institute, National Institutes of Health. We would like to thank the NCI CCR Frederick Flow Cytometry Core, Optical Microscopy and Image Analysis Lab (OMAL), and the Laboratory Animal Sciences Program for their assistance in access to the instrumentation used to perform the experiments in this manuscript. We would like to thank the CCR Volume Electron Microscopy (CVEM) core, particularly Dr. Narayan and Dr. Harned, for their help in sample preparation and data collection in SEM.

