## Supporting Information for "Modulating Neutrophil Extracellular Trap Formation *In Vivo* with Locoregional Precision using Differently Charged Self-Assembled Hydrogels"

### Materials and Methods

#### Safety Statement

No unexpected or unusually high safety hazards were encountered during the development of this project.

#### Peptide Synthesis

MAX8, HLT2, TLK5, and TLE5 were synthesized using a CEM Liberty Blue microwave-assisted peptide synthesizer on ProTide Rink amide resin (0.58-0.65 mmol/g) at 0.25 mmol scale. Fmoc deprotection cycles were performed with 20% piperidine in DMF (15s at 75°C followed by 50s at 90°C). The first 11 couplings (residues 20-10) were performed at 90°C for 4 min using Fmoc-protected amino acids (5 eq, 0.2 M in DMF), Oxyma (5 eq, 1 M in DMF), and DIC (5 eq, 0.5 M in DMF). The Valine after the ᴅ-Proline amino acid was triple coupled with HCTU (5 eq, 0.45 M in DMF) and DIPEA (10 eq, 2M in NMP) for 30 min at room temperature. The same coupling conditions were used for the subsequent two residues with double coupling. The remaining six residues were coupled at 90°C as described above. MAX8 and HLT2 have a free N-termini. For TLK5 and TLE5, the N-terminus was acetylated with acetic anhydride (10% v/v in DMF). Resin was washed with DCM and DMF, dried under vacuum before cleavage. Peptides were cleaved with 25 mL of 95% Trifluoroacetic acid (TFA), 2.5% Triisopropylsilane, and 2.5% water for 3 h. TFA was removed by argon gas flushing, and crude peptides were precipitated with cold diethyl ether followed by two washes. Crude peptides were dissolved in water and lyophilized before mass confirmation and purification.

#### Peptide Purification

Peptides MAX8, HLT2, TLK5 were purified by preparative RP-HPLC on a Waters 600 system equipped with a Waters 2489 or 2487 UV/Vis Detector using a Vydac 218TP C18 column (250 x 4.6 mm, 5 µm) at a flow rate of 8.0 mL/min and a column temperature of 40°C. The UV trace was monitored at 220 nm. Gradients and solvent systems for preparative RP-HPLC were:

*MAX8*: 100% *Standard A (0.1% TFA in* $H_2O$*)* for 2 min followed by a linear gradient from 0 to 21% *Standard B (90% Acetonitrile, 9.9%* $H_2O$*, 0.1% TFA)* in 5 min (4.2% Standard B/min). Then a linear gradient of 0.5% Standard B/min for 40 min. MAX8 elutes at 36% Standard B.

*HLT2:* 100% Standard A for 2 min followed by a linear gradient from 0 to 19% Standard B in 5 min (3.8% Standard B/min). Then a linear gradient of 0.5% Standard B/min for 40 min. HLT2 elutes at 34% Standard B.

*TLK5:* 100% Standard A for 2 min followed by a linear gradient from 0 to 23% Standard B in 5 min (4.6% Standard B/min). Then a linear gradient of 0.5% Standard B/min for 40 min. TLK5 elutes at 38% Standard B.

Peptide purity was assessed by LCMS (Shimadzu LCMS 2020) using a Phenomenex Luna C18 column 100 Å (150 x 3 mm, 5 µm) with a linear gradient from 0 to 90% Standard B2 (0.1% Formic acid in acetonitrile) at 2% B2/min and a column temperature of 40°C. Purity was also assessed by Analytical RP-HPLC (Agilent 1200 series) using a Vydac 218TP C18 column (250 x 4.6 mm, 5 µm) at 40°C with a linear gradient from 0% to 100% Standard B at a rate of 1% B/min. Peptides were lyophilized to afford a white powder.

*TLE5* was purified by preparative RP-HPLC using a Water 600 system with 2489 UV/vis detector in a Phenomenex PolymerX RP-1 column (250 x 21.2 mm, 10 µm, heated to 40°C at 8 mL/min. The gradient was: 100% Standard C (20 mM $NH_4HCO_3$) for 2 min followed by a linear gradient from 0 to 16% Standard D (80% Acetonitrile, 20% 20 mM $NH_4HCO_3$) in 5 min (3.2% Standard D/min). Then a linear gradient of 0.5% Standard D/min for 40 min. TLE5 eluted at 31% Standard D. Purity was assessed by LCMS and analytical HPLC as previously described but with a Phenomenex PolymerX RP-1 column (250 x 4.6 mm, 10 µm) using gradients of 1% Standard D/min. After lyophilization, TLE5 was converted to sodium salt by dissolving the peptide in water and adjusting the pH to 7.4 with addition of aqueous NaOH. The peptide solution was frozen and lyophilized again to afford a white powder.

All peptides were dissolved in ultrapure or endotoxin free water and sterile filtered with a 0.2 µm polystyrene filter before being lyophilized again for their use in further experiments.

**Peptide Characterization**

Peptides MAX8 and HLT2 have been characterized and published previously.[1, 2] TLK5 and TLE5 are new designs and were fully characterized for their secondary structure, self-assembly, gelation, and viscoelastic properties as described below.

***Circular dichroism (CD) spectroscopy***

Characterization of the secondary structure and β-sheet formation as a function of buffer and temperature was performed using a Jasco J-1500 CD spectrometer. Ice-cold peptide stock solutions in ultrapure water at 300 µM or 2 wt.% (8.03 mM TLK5, 8.76 mM TLE5) were mixed with ice-cold 2X HEPES Buffer solution (HBS, 50 mM HEPES, 300 mM NaCl, pH 7.4) for a final concentration of 150 µM or 1 wt.% (4.02 mM TLK5, 4.38 mM TLE5). The samples were analyzed in a 1 mm (for 150 µM) or 0.1 mm (for 1 wt.%) high precision cuvette (Hellma Analytics). The samples were allowed to equilibrate on the spectrometer for 10 min at 2 °C, and then wavelength scans from 260 to 200 nm were taken every 5 °C. Between every temperature interval, the samples were allowed to equilibrate for 10 min before the measurement. Data were converted to Mean Residue Ellipticity (MRE) calculated as:

MRE= $\theta obs \,/\, (10\, l\, c\, r)$

where $\theta_{obs}$ is the ellipticity in millidegrees, l is the light path on the cuvette, c is the molar concentration (M), and r is the number of peptide bonds (residues) in the peptide.

***Transmission Electron microscopy (TEM)***

1 wt.% peptide gels were prepared by mixing 2 wt.% peptide solutions in ultrapure water with 2X HBS (1:1). Samples were cured overnight at 37 °C.  An aliquot of peptide gel was diluted 40-fold for TLK5 and 100-fold for TLE5 in ultrapure water and mixed thoroughly. 3.5 µL of the diluted gel were deposited on top of a glow discharge-treated grid (400-square mesh, CF400-CU, Electron Microscopy Science, Inc., PELCO easiGlow, Ted Pella, Inc.) and allowed to sit for 1 min. The grid was blotted with clean filter paper and washed three times with ultrapure water by adding a 3.5 µL water droplet on top, followed by blotting with paper. Samples were negatively stained with 3.5 µL of 2 wt.% uranyl acetate in water. Samples were dried overnight at room temperature before imaging on a FEI Tecnai 12 Twin transmission electron microscope ay 80 kV acceleration voltage. Images were captures with an AMT XR16 charge-coupled device (CDC) bottom-mount camera (Advanced Microscopy Techniques, Corp.). Fibril width was measured using ImageJ (NIH).

***Oscillatory Rheology***

Characterization of gelation and viscoelastic properties of the peptide gels was performed using a Discovery HR20 (TA Instruments) using a 25 mm stainless steel parallel plate geometry and 500 µm gap height. Peptides were dissolved in ice-cold ultrapure water (20 mg/mL) and diluted 1:1 with ice-cold 2X HBS immediately before the test. Then, 300 µL of 1wt.% (10 mg/mL) peptide solution was immediately placed on the cold rheometer stage at 5 °C. Geometry was lowered to the measurement gap and covered with mineral oil (ASTM oil standard) to prevent drying. Sample was allowed to soak at 5 °C for 10 s followed by a temperature ramp to 37 °C (17.45 °C/min) at a strain of 0.2% and angular frequency of 6.0 rad/s. The evolution of the storage (G') and loss (G'') moduli were monitored for 1 h with an oscillatory time sweep experiment at 37 °C, 0.2% strain, and 6.0 rad/s angular frequency. Then, 1000% strain was applied for 30 s followed by another oscillation time sweep at 0.2% to evaluate the thixotropic properties and recovery of the peptide gels. To evaluate the time dependence of the viscoelastic properties, oscillation frequency sweeps were performed on the same sample at 37 °C, 0.2% strain, and a frequency range from 0.1 to 100 rad/s. Then, oscillation amplitude seeps were performed at 37 °C, 6.0 rad/s angular frequency, and a strain range from 0.1 to 1000% to determine the viscoelastic behavior and flow point of each gel. To estimate the G' stabilization time, the first derivative of G' as a function of time was calculated using GraphPad Prism. The stabilization time was determined at the time where there is ≤ 10% change in the highest first derivative value.

**Peptide gelation for *in vivo* experiments**

After purification, peptides were dissolved in endotoxin-free water, sterile-filtered through a 0.22 µm PVDF Durapore Membrane (Millex®-GV Merck Millipore, Tullagreen Irl.), and filtered through a Mustang™ E 0.2 µm, 25 mm Acrodisc® unit under asceptic conditions. Peptides were then lyophilized and stored at -20°C until use. Sterile-filtered peptide powders were dissolved in cold ultrapure water under sterile conditions at 2 wt.% (20 mg/mL). After complete peptide dissolution, pre-chilled sterile and endotoxin free 2X HBS (50 mM HEPES, 300 mM NaCl, pH 7.4) was added to the peptide solution in a 1:1 ratio. The final concentration of the hydrogel was 1% wt. (10 mg/mL) in 25 mM HEPES, 150 mM NaCl, pH 7.4. Peptide solutions were loaded into sterile syringes, placed on sterile pouches, and transfer to an incubator at 37 °C overnight to promote complete gelation. Endotoxin levels were quantified for each gel using the ToxinSensor™ Chromogenic LAL Endotoxin Assay, Endotoxin Detection System (GenScript). No endotoxins were detected for HLT2 and MAX8, and TLK5 and TLE% had levels below or equal to 0.1 EU/mL, that is the limit of detection for this protocol. Osmolarity was measured using a Vapor Pressure Osmometer (VAPRO, Wescor). The osmolarity was within a range of 353 – 387 mmol/kg.

***In vivo* Subcutaneous and intramuscular injection model**

All experimental procedures were approved by the NCI Animal Care and Use Committee (ACUC) and performed according to the Animal Welfare Act and NIH guidelines for the care and use of laboratory animals. Female C57BL/6J mice (8 to 12 weeks old) were obtained from The Jackson Laboratories. Animals were anesthetized in an induction chamber using 3-4% Isoflurane (1-1.5 L/min oxygen flow rate). Then animals were placed in a surgery board at 37 °C with eye lubricant and maintained anesthesia at 2% isoflurane. Hair from the back or leg was removed using clippers, followed by the application of facial hair removal cream. The cream was removed, and the area was neutralized with 1 % (v/v) acetic acid solution to prevent irritation. The mouse skin was cleaned with alcohol swabs before injection. Hydrogels were injected into the subcutaneous space of their dorsal flank using a 27-gauge needle or into the gastrocnemius muscle, and the animals were allowed to recover in a heating pad with food, water, and/or DietGel recovery. At specific time points during the study, animals were anesthetized with 4-5 % Isoflurane and euthanized by $CO_2$ asphyxiation, followed by a secondary method of euthanasia before tissue harvesting.

For histological analysis, mice received two 150 µL injections to the left and right sides of their dorsal flanks. For intramuscular injections, mice received a 50 µL injection in the right gastrocnemius muscle. Animals were euthanized at 3-, 7-, 14-, and 30-days post-injection. For flow cytometry and cytokine-chemokine quantification, mice received four 100 µL injections. For NET modulation studies, mice received four 100 µL injections, and implants from the same mouse were used for both histological analysis and cytokine/chemokine studies. For degradation studies, mice received one 150 µL injection in the right flank. For locoregional control studies, TLE5 peptide gel was drawn first into the syringe to a final volume of 50 µL. Then, TLK5 was drawn next carefully to prevent bubbles to a final volume of 50 µL, resulting in a single syringe with two different materials as shown in Figure 7a. Hydrogels were injected slowly into the subcutaneous space to prevent mixing of both materials *in vivo*.

**Degradation studies using ultrasound imaging**

150 µL of peptide gel was subcutaneously administered on the lower back of 6–8 weeks old C57BL/6J mice (n = 4). Hydrogel implant degradation was evaluated by B-mode ultrasound imaging using the Vevo2100 preclinical scanner (VisualSonics Inc., Toronto, CA) at 1-, 2-, 3-, 4-, 5-, 10-, 12-, 15-, 17-, 19-, 24-, 31-, and 38- days post-injection. Ultrasound images were acquired with a MS 550S (40 MHz) linear array transducer with a step size of 0.076 mm. Hydrogel volumes were measured for each time point using the parallel contour algorithm (Vevo LAB software ver. 1.7.1, Visual Sonics Inc., Toronto, CA).

**Histological analysis and immunofluorescence staining**

After tissue harvesting, implanted hydrogels on the skin were fixed with 10% neutral buffered formalin overnight. The skin and implants were trimmed, cut, and sent for processing, sectioning, and staining to Histoserv, Inc. (Germantown, MD). Hematoxylin & eosin and Masson's trichome stained slides were imaged using an Aperio Slide Scan (40X) and deposited to Indica Labs HALO Link™ by the Molecular Histopathology Laboratory at the NCI Frederick.

5 µm tissue slides were deparaffinized with two xylenes treatments for 5 min each, followed by rehydration with a series EtOH treatments (from 100% to 50% EtOH in DI water). Antigens were retrieved by boiling in sodium citrate buffer (10mM sodium citrate, 0.05% Tween 20 at pH 6.0).

Then, slides were rinsed and permeabilized twice with 0.1% PBST (0.1% Tween 20 in 1X PBS pH 7.4) for 2 min and blocked with 1% w/v BSA in PBS for 30 min. Tissue sections were incubated with primary antibodies (**Table S1**) overnight at 4 °C, then rinsed with PBST twice and incubated with secondary antibodies (**Table S1**) for 1h at room temperature. Tissue slides were rinsed and incubated with SYTOX Blue for 30 min before a last wash session and mounting with ProLong® Diamond Antifade reagent (Invitrogen). Slides were analyzed using a Zeiss LSM780 Laser scanning confocal microscope at the Optical Microscopy and Analysis Laboratory at the NCI Frederick.

***Table S1.*** *Primary and secondary antibodies for immunofluorescence staining*

| Specificity | Reactivity | Fluorochrome | Isotype | Clone | Vendor Cat # | Concentration | Dilution |
|---|---|---|---|---|---|---|---|
| *Primary Antibodies* | | | | | | | |
| Ly6G | Mouse | - | Rat IgG2a, κ | 1A8 | Biolegend 127602 | 0.5 mg/mL | 1:500 |
| Neutrophil Elastase | Mouse, Human | - | Rabbit IgG | JF098-6 | Novus NBP2-66972 | 1 mg/mL | 1:200 |
| Myeloperoxidase | Mouse Human | - | Goat IgG | Polyclonal | R&D systems AF3667 | 0.2 mg/mL | 1:100 |
| SYTOX™ Blue (DNA) | - | SYTOX™ Blue | - | - | Thermo Fisher Scientific S11348 | 5 mM | 1:1000 |
| *Secondary Antibodies* | | | | | | | |
| Goat IgG | Goat | Alexa Fluor™ 647 | Donkey IgG | Polyclonal | Thermo Fisher Scientific A21447 | 2 mg/mL | 1:500 |
| Rat IgG | Rat | Alexa Fluor™ 568 | Donkey IgG | Polyclonal | Thermo Fisher Scientific A78946 | 2 mg/mL | 1:500 |
| Rabbit IgG | Rabbit | Alexa Fluor™ 488 | Donkey IgG | Polyclonal | Thermo Fisher Scientific A21206 | 2 mg/mL | 1:500 |

### Flow cytometry and immunophenotyping

Hydrogel implants were harvested after 3-, 7-, and 14-days post-injection (4 implants per animal) with any connective and fat tissue trimmed. Then, implants were cut with a scalpel, and digested with 5 mL of Liberase™ (Thermolysin Medium, 0.2 WU/mL) and 80 U/mL DNase in 1X PBS at 37°C for 45 min with gentle agitation. After digestion, the cell suspension was filtered through a 40 µm cell filter. The filter was rinsed several times with 1% BSA 2 mM EDTA in DPBS to a final volume of 20 mL. Cell suspensions were centrifuged for 5 min at 1200 rpm at 4°C, washed with 1X PBS once, centrifuged, and resuspended in 1X PBS. Cells were counted, concentration adjusted, and 1 x $10^6$ cells were placed in each tube for staining (**Table S2**). Cells were incubated with 100 µL of fixable live/dead Zombie NIR (1:2000 dilution) in PBS for 20 min on ice, followed by two washes with Stained buffer (FBS in DPBS, BD Biosciences). Then, cells were blocked with Mouse BD Fc block™ CD16/CD32 for 5 min at 4°C before adding the antibody cocktail (**Table S2**) to a final volume of 100 µL. Cells were incubated on ice for 30 min, washed twice with stain buffer, fixed with BD Cytofix™ fixation buffer (200 µL for 15 min on ice) while being filtered through a 30 µm cell trainer, followed by a last wash. Cells were resuspended in stain buffer and analyzed using a Cytek Aurora Spectral Flow Cytometer and data was analyzed on SpectroFlo ®. Gates were determined using Fluorescent minus one (FMO) controls.

***Table S2.*** *Flow cytometry panel for identifying infiltrating cells*

| Specificity | Fluorochrome | Clone | Isotype | Vendor | Catalog # | Titer (ng/test) |
|---|---|---|---|---|---|---|
| Viability | Zombie NIR | | | Biolegend | 423106 | 100 µL of 1:2000 dilution from stock |
| CD45 | BV480 | 30-F11 | Rat LOU/M IgG2b, κ | BD | 566168 | 100 |
| CD11b | BV570 | M1/70 | Rat IgG2b, κ | Biolegend | 101233 | 200 |
| Ly6G | PerCP-Cy5.5 | 1A8 | Rat Lewis IgG2a, κ | BD | 560602 | 400 |
| F4/80 | BUV737 | T45-2342 | Rat WI IgG2a, κ | BD | 749283 | 400 |
| Ly6C | Dylight350 | ER-MP20 | Rat IgG2a | Novus Biologicals | NB100-65413UV | 710 |

**Scanning Electron Microscopy of implants and native gels**

SEM sample preparation and imaging was performed by the Center for Cancer Research Volume Electron Microscopy (CvEM) core at the NCI. Peptide implants were harvested from the mice as described previously at days 3 and 28 post-injection. Implants were trimmed and cleaned, followed by fixation with a solution of 4% formaldehyde and 2% glutaraldehyde (v/v) in 0.1 M cacodylate buffer, pH 7.4. Implants were subsequently stained and fixed using a 1% osmium tetroxide solution and broken to reveal the periphery area of the implants where NETs are typically observed by histology. Then, samples were dehydrated using a standard alcohol dehydration process ranging from 35% to 100% with final dehydration completed using Tetramethylsilane. The dried samples were coated with an iridium layer of approximately 5 nm using an Emitech K575X sputter coater. Samples were imaged with a ZEISS GeminiSEM 450. The same sample preparation procedure was performed in TLK5 and TLE5 native hydrogels to observe the morphology of the materials before injection and *in vivo* modifications.

**Multiplex immunoassay for cytokine and chemokine quantification**

Peptide hydrogels (1 wt.%, 10 mg/mL) were prepared under sterile conditions as described before. Mice received four 100 µL peptide gel injections at the dorsal aspect (n = 5 mice per time point). At 1-, 3-, 7-, and 14-days post-injection, mice were euthanized according to the ACUC approved protocol. Hydrogel implants (4 implants) were retrieved from the subcutaneous space, trimmed, rinsed with cold PBS, and placed on 1 mL of 1X Halt™ protease inhibitor cocktail (Thermo Scientific) in ice cold PBS. Implants were cut into small pieces and homogenized before adding Triton X-100 (IBI scientific) and homogenized again. Proteins were extracted from the samples by freeze-thaw lysis by fast freezing samples in liquid Nitrogen, follow by thawing on ice and pulsed vortexing. Lysates were centrifuged at 10,000 xg for 5 min at 4°C, and supernatants were aliquoted for storage at -80°C or protein quantification. Protein concentration as determined with the Pierce™ BCA Protein Assay Kit (Thermo Scientific) in 1:10 diluted samples with a working range = 20-2,000 µg/mL as specified by the manufactured protocol.

Cytokines and chemokines were quantified using a custom mouse panel LEGENplex™ bead-based multiplex immunoassay kit for CXCL1 (KC), CCL3 (MIP-1α), CCL5 (RANTES), IL-1β, IFN-γ, IL-6, G-CSF, TNF-α, IFN-β, CCL2 (MCP-1), and GM-CSF. Samples were prepared according to the manufacturer's instructions. Briefly, lysates were diluted 1:2 with assay buffer prior to quantification. All samples and reagents were warmed to room temperature and run in duplicate. 25 µL of assay buffer and sample or standard were added to a V-bottom plate with 25 µL of mixed beads. Samples were incubated for 2h at room temperature under constant shaking, followed by a wash step before incubation with biotinylated detection antibodies for 1h at room temperature under shaking. Then, 25 µL of SA-PE were added to each sample without washing and incubated for 30 min at room temperature under constant shaking. Samples were washed, transfered to 5 mL Corning Falcon round-bottom polystyrene tubes and analyzed in a Cytek Aurora spectral flow cytometer. Data was analyzed using Biolegend's LEGENDplex™ data analysis software.

ELISA assays were used to quantify Neutrophil Elastase, Myeloperoxidase, and G-CSF from the implant lysates. DuoSet™ ELISA development systems for Mouse Neutrophil Elastase/ELA2 (DY4517), Mouse Myeloperoxidase (DY3667), and Mouse G-CSF (DY414) (R&D Systems, Inc, Minneapolis, MN) were used according to the manufactured protocol. Briefly, 96 well microplates were coated overnight at room temperature with capture antibody at the working concentration (NE: 1 µg/mL, MPO: 800 ng/mL, G-CSF: 2 µg/mL). Then, the plates were washed three times with 1X wash buffer and blocked with 1% BSA in PBS reagent diluent for at least 1 h. After washing the blocking solution three times, sample and standards were added to each well and incubated for 2 h at room temperature. Wells were washed three times before adding the detection antibody at the working concentration (NE: 100 ng/mL, MPO: 20 ng/mL, G-CSF: 200 ng/mL) and incubated for 2 h at room temperature. Then, plates were washed again and Streptavidin-HRP (1:200 dilution) was added for 20 min limited from light, followed by another washing process and addition of the substrate solution for 20 min. Reaction was stopped with 2N sulfuric acid and analyzed in a BioTek Synergy neo 2 multi-mode plate reader set at 450 nm with correction at 540 nm.

**Statistics**

Treatment groups for each experiment were created by randomized mouse selection upon receipt at the facility. For comparisons between groups at specific timepoints, ordinary one-way analysis of variance (ANOVA) with Tukey's correction for multiple comparisons were performed. Normality tests were not performed due to the small sample sizes. The sample size for each experiment is stated in the figure caption and range from an n = 3 to 8. Data collection and analysis were not performed blind to the conditions of the experiments. P values are shown in the figure captions and SI.

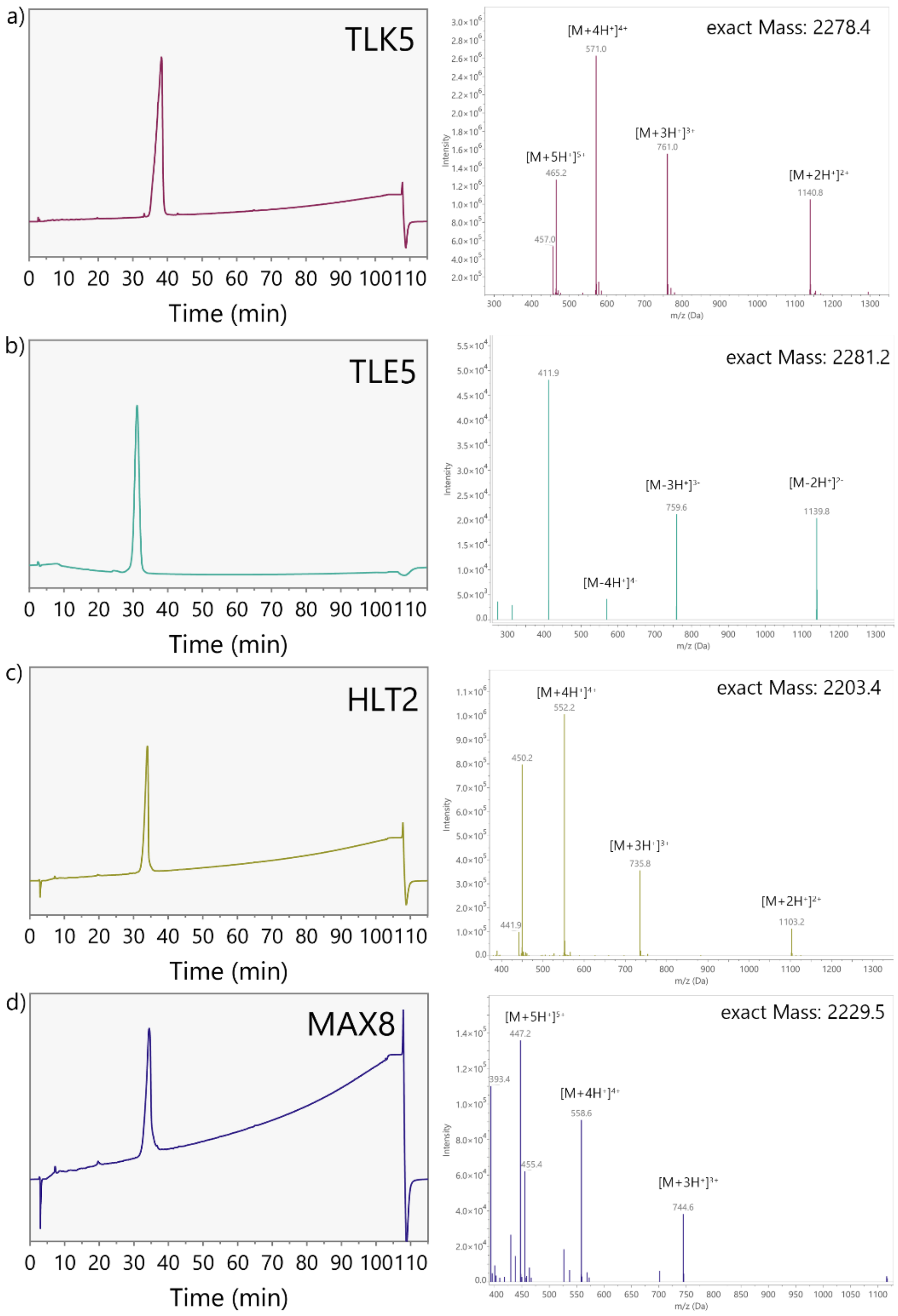

***Figure S1.*** RP-HPLC trace and ESI mass spectra of a) TLK5, tR = 38 min, calculated mass: $[M+H^+]^{1+}$ = 2279.4, $[M+2H^+]^{2+}$ = 1140.2, $[M+3H^+]^{3+}$ = 760.5, $[M+4H^+]^{4}$ = 570.6, $[M+5H^+]^{5+}$ = 456.7. b) TLE5, tR = 31 min, calculated mass: $[M-H^+]^{1-}$ = 2280.2, $[M-2H^+]^{2-}$ = 1139.6, $[M-3H^+]^{3-}$ = 759.4, $[M-4H^+]^{4-}$ = 569.3, $[M-5H^+]^{5-}$ = 455.2. c) HLT2, tR = 34 min, calculated mass: $[M+H^+]^{1+}$ = 2204.4, $[M+2H^+]^{2+}$ = 1102.7, $[M+3H^+]^{3+}$ = 735.5, $[M+4H^+]^{4}$ = 551.9, $[M+5H^+]^{5+}$ = 441.7. d) MAX8, tR = 36 min, calculated mass: $[M+H^+]^{1+}$ = 2230.5, $[M+2H^+]^{2+}$ = 1115.8, $[M+3H^+]^{3+}$ = 744.2, $[M+4H^+]^{4}$ = 558.4, $[M+5H^+]^{5+}$ = 446.9.

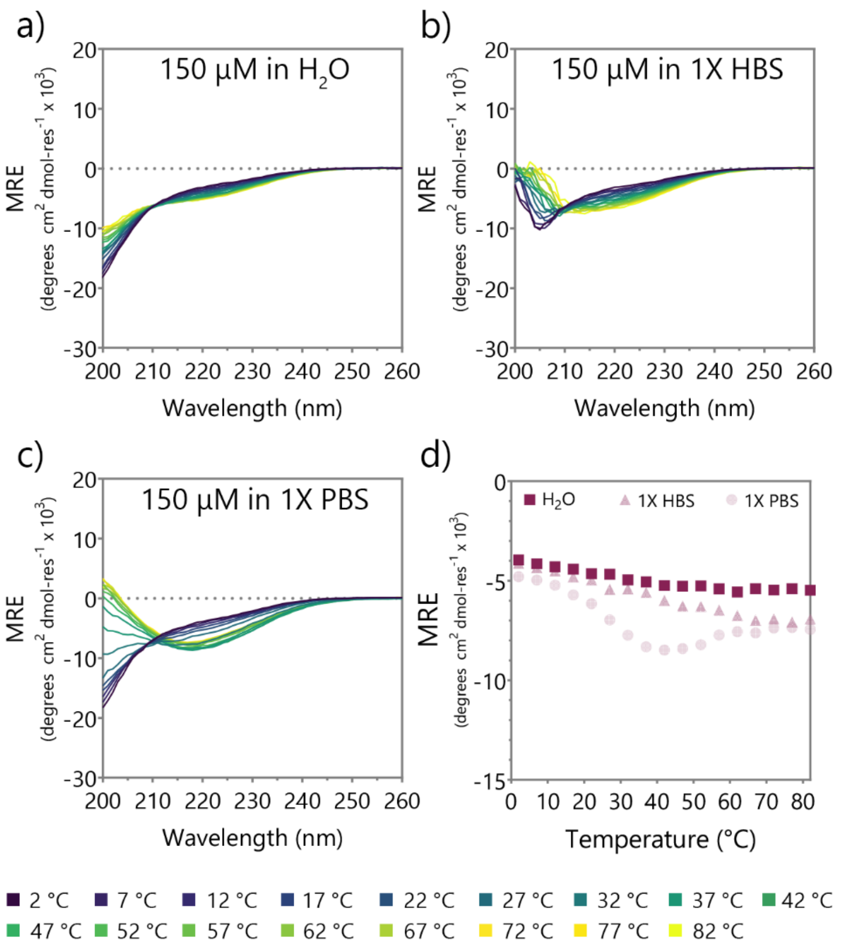

***Figure S2.*** Temperature-dependent CD spectra of TLK5 at 150 µM in a) water, b) 1X HBS pH 7.4, and c) 1X PBS pH 7.4. d) MRE values at 216 nm as a function of temperature to monitor β-sheet formation. At this concentration, TLK5 does not form a β-sheet and assemble. In 1X PBS, TLK5 forms a β-sheet with the characteristic minimum at 216 nm but precipitates out of the solution.

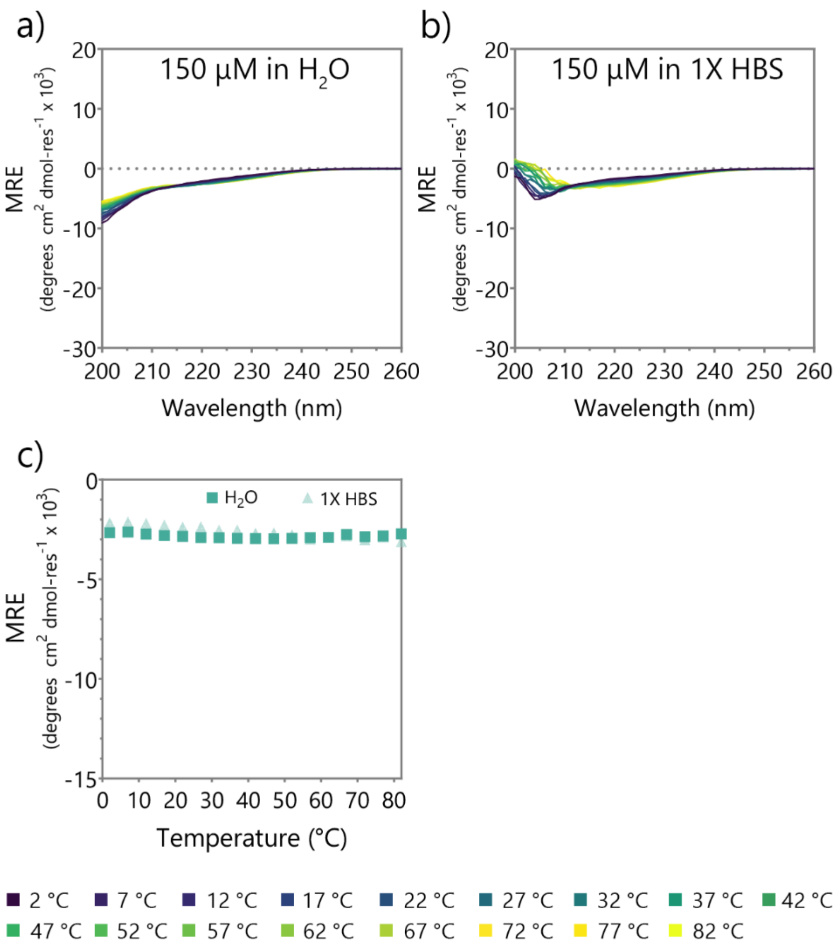

***Figure S3.*** Temperature-dependent CD spectra of TLE5 at 150 µM in a) water and b) 1X HBS pH 7.4. d) MRE values at 216 nm as a function of temperature to monitor β-sheet formation. At this concentration, TLE5 does not form a β-sheet and assemble.

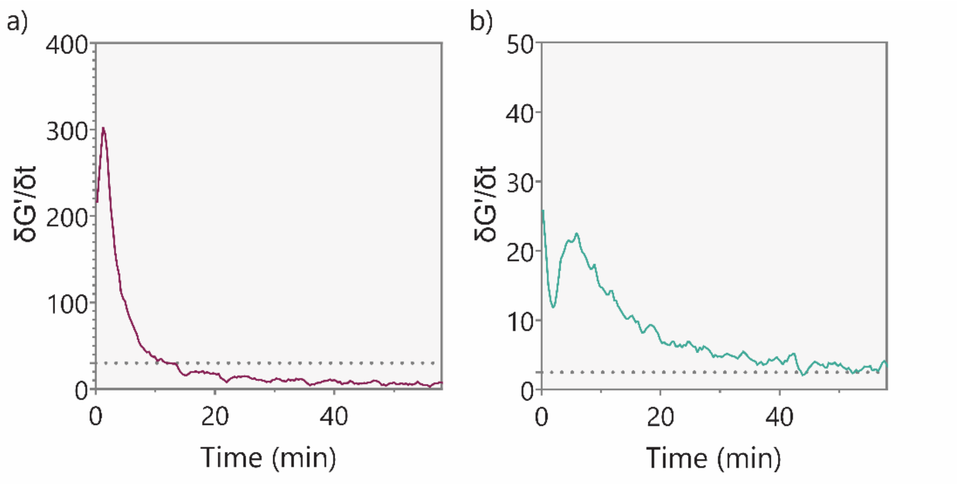

***Figure S4.*** Time for gel stabilization. First derivative of the averaged storage modulus vs time represents the change in G’ as a function of time. Tiem of stabilization was determined as <= 10% of change in G’.

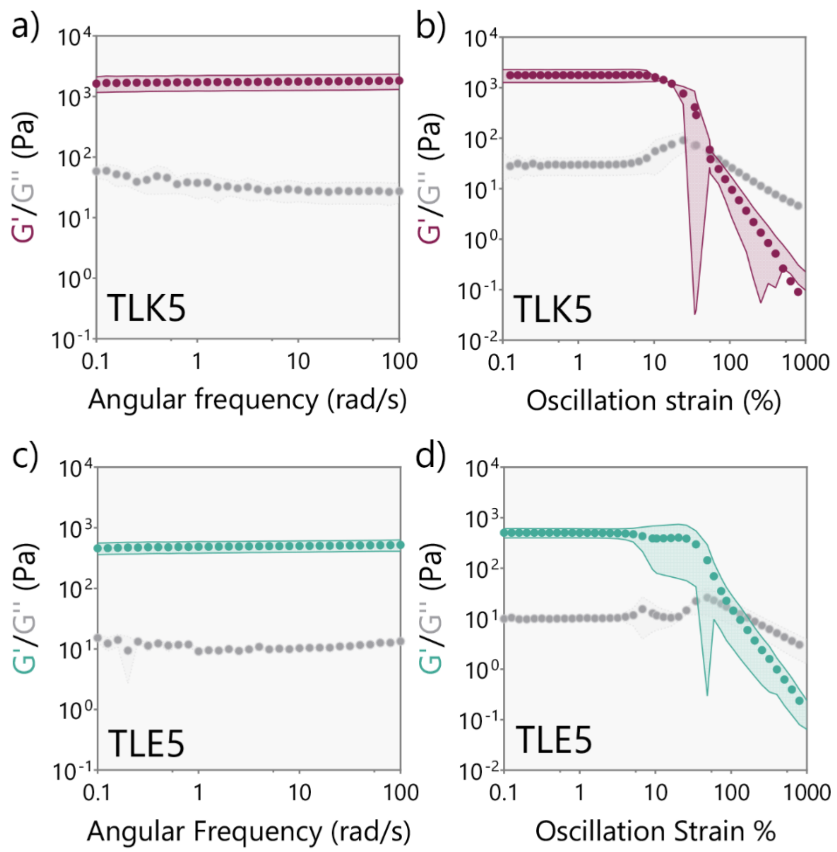

***Figure S5.*** Rheological characterization of peptide gels. Frequency sweep (a) and amplitude sweep (b) of TLK5 1 wt.% gel. Frequency sweep (c) and amplitude sweep (d) of TLE5 1 wt.% gel. Data are shown as mean and standard deviation (with error bands) n = 3.

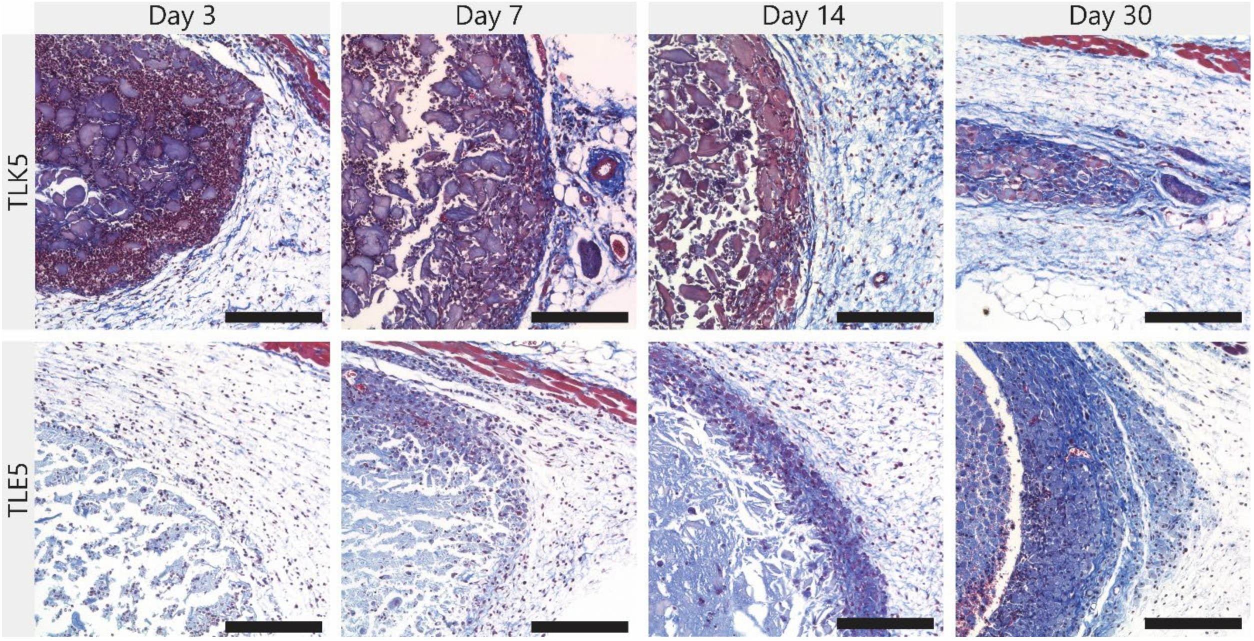

***Figure S6.*** Tissue sections of the different hydrogel implants in the subcutaneous space stained with Masson's Trichrome at different time points. Scale bar = 200 µm.

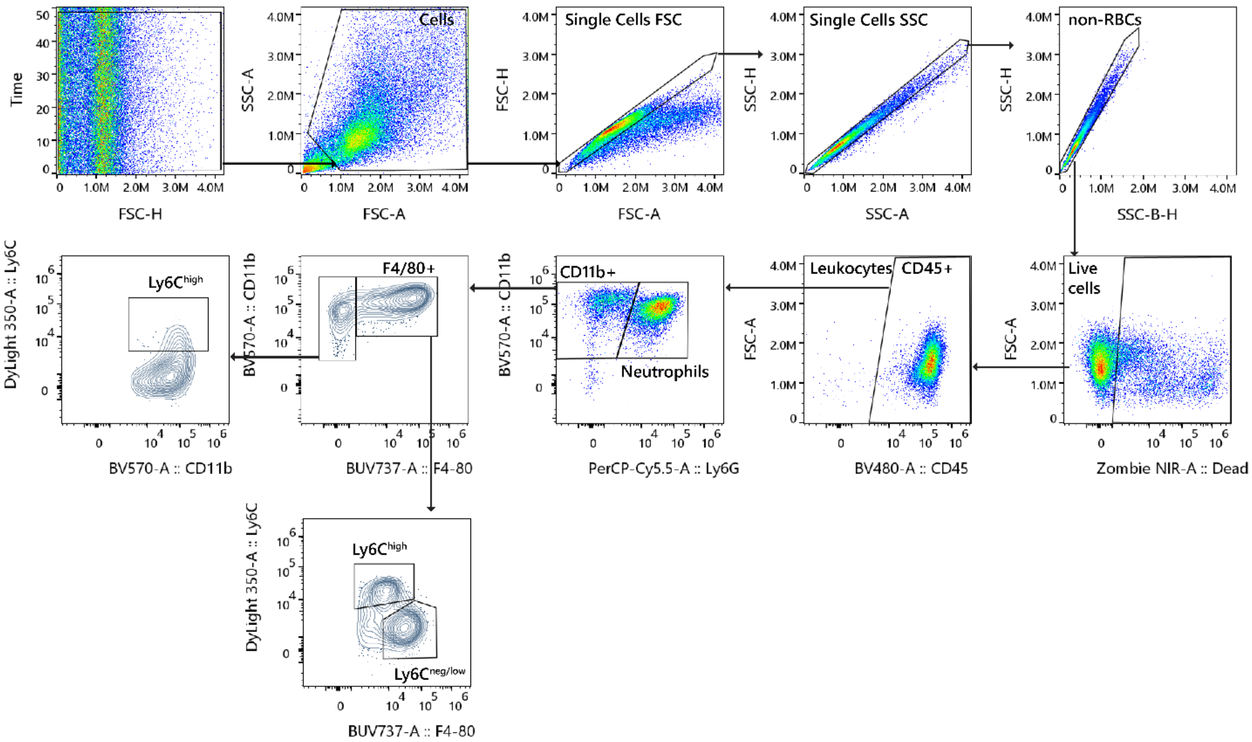

***Figure S7.*** Gating strategy for myeloid panel to identify Neutrophils (CD45+CD11b+Ly6G+), and macrophages (CD45+CD11b+F4/80+Ly6Chigh or low/neg). Data was cleaned by gating on steady flow, viable cells, singlets, non-red blood cells, and live cells.

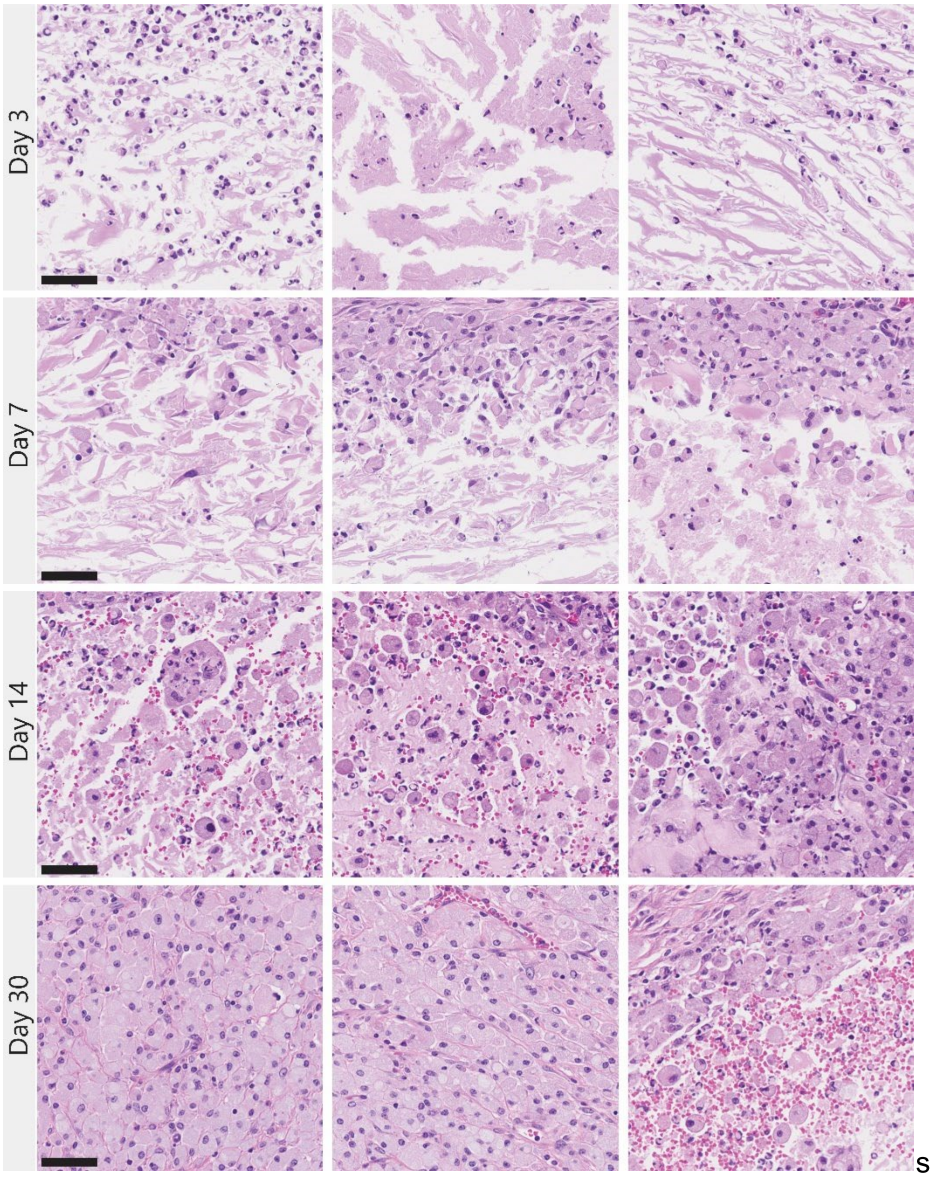

***Figure S9.*** H&E-stained tissue sections of TLE5 implants. Hydrogel presents a lower degree of cellular infiltration, mostly in the periphery and surface of the implants. At later time points, there is more dense cell infiltration in the periphery and the core contains red blood cells, neutrophils, and macrophages. Scale bar = 50 µm.

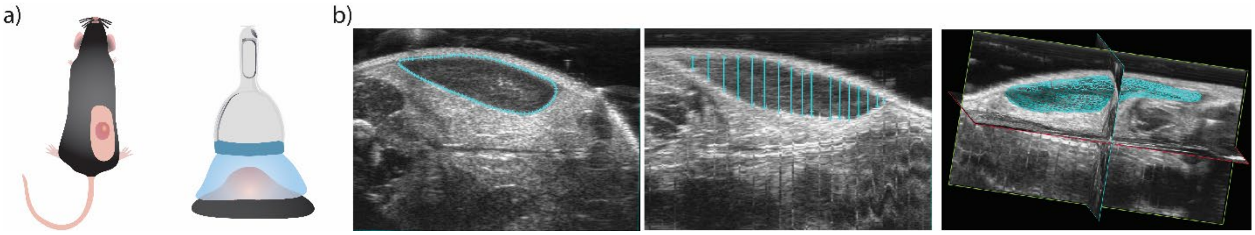

***Figure S10.*** Monitoring hydrogel degradation using ultrasound imaging. a) Mice were subcutaneously injected with 150 μL hydrogels in the dorsal flank and the implant volume was obtained with ultrasound imaging. b) Examples of ultrasound images and volume determination using Vevo LAB software.

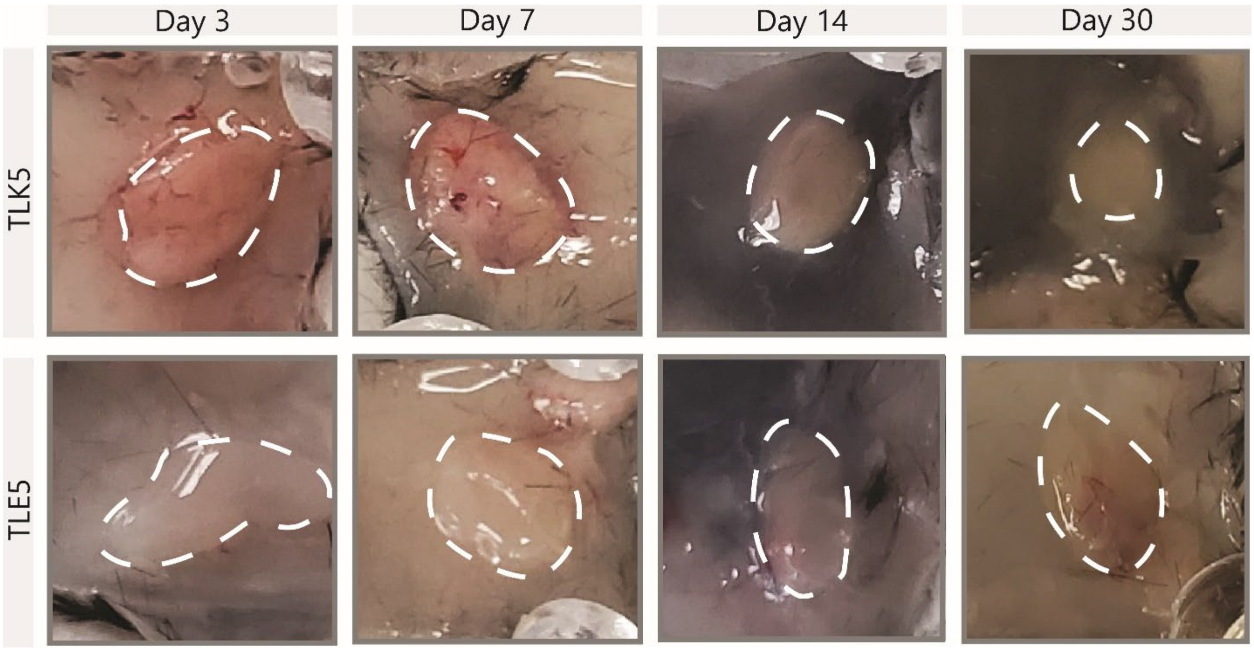

***Figure S11.*** Gross histology of subcutaneous TLK5 and TLE5 implants over time.

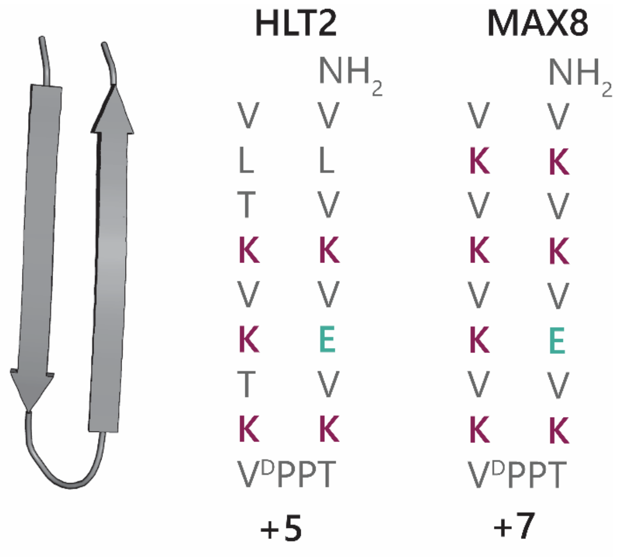

***Figure S12.*** Peptide sequences of positively charged gel-forming peptides HLT2 and MAX8.

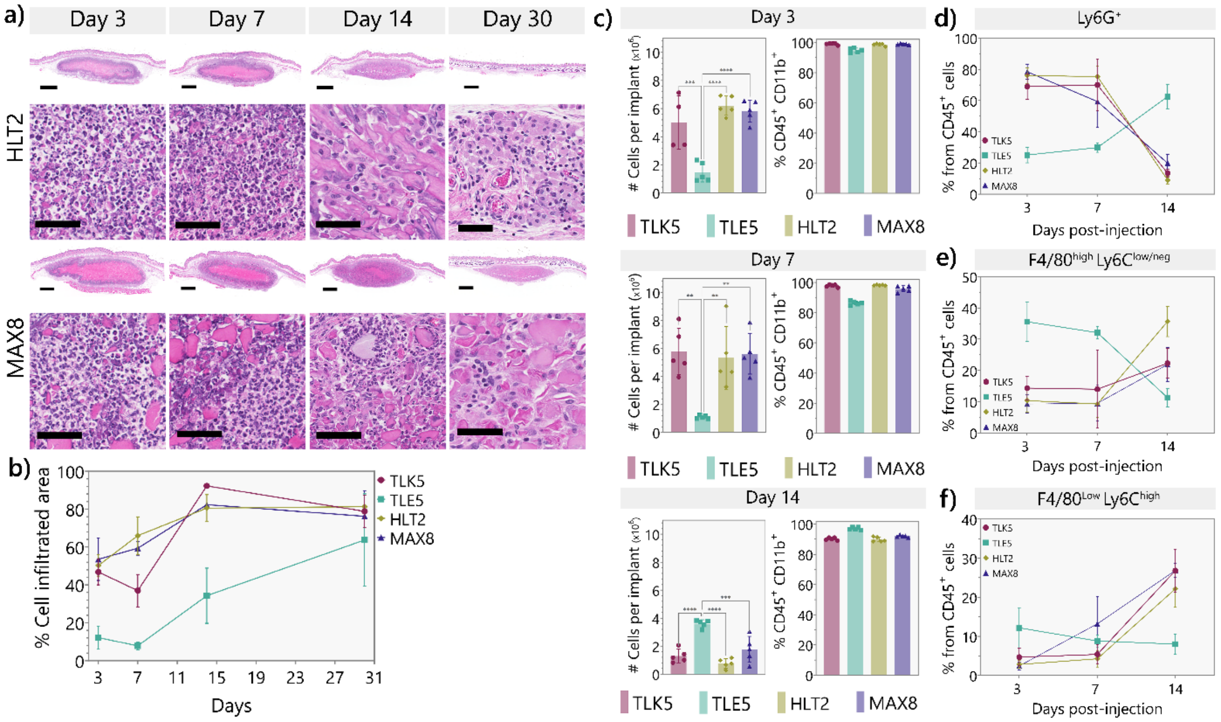

***Figure S13.*** Characterization of the immune response to HLT2 and MAX8. **a)** H&E-stained tissue sections of implants for HLT2 and MAX8 gels at 3-, 7-, 14-, and 30- days post-injection. **b)** Percentage of infiltrated area over time determined by histology, n = 3 mice. *Statistical differences at day 3: TLE5 vs TLK5, HLT2 ** p-value < 0.003, TLE5 vs MAX8 *** p-value 0.0008; at day 7: TLE5 vs TLK5 ** p-value 0.0037, TLE5 vs MAX8, HLT2 **** p-value <0.0001, at day 14: TLE5 vs HLT2, MAX8 ** p-value < 0.003, TLE5 vs TLK5 *** p-value 0.0004.* **c)** Number of cells recovered per implant and % myeloid cells ($CD45^+CD11b^+$) at day 3, day 7, and day 14 post-injection. n = 5 mice. **d)** Percentage of $CD45^+CD11b^+Ly6G^+$ cells (Neutrophils) from total leukocytes for each peptide hydrogel at different timepoints. n = 5 mice. *Statistical comparison: day 3 TLK5, HLT2, MAX8 vs. TLE5 p-value < 0.0001; day 7 TLK5, HLT2 vs. TLE5 p-value < 0.0005, TLE5 vs. MAX8 p-value 0.0075; day 14 TLK5, HLT2, MAX8 vs. TLE5 p-value <0.0001, HLT2 vs. MAX8 p-value 0.0193.* **e)** Percentage of $CD45^+CD11b^+Ly6G^{neg}F4/80^{high}Ly6C^{low/neg}$ cells from total $CD45^+$ leukocytes for each peptide hydrogel at different timepoints. *Statistical comparison: Day 3, TLK5, HLT2, MAX8 vs. TLE5 p-value < 0.0001. Day 7, TLK5 vs. TLE5 p-value 0.0057, TLE5 vs. HLT2 and MAX8 p-value < 0.0008. Day 14 TLK5 vs. TLE5 and HLT2 p-value <0.0080, TLE5 vs. HLT2 p-value < 0.0001, TLE5 vs MAX8 p-value 0.0103, HLT2 vs. MAX8 p-value 0.0012.* **f)** Percentage of $CD45^+CD11b^+Ly6G^{neg}F4/80^{low}Ly6C^{high}$ cells (monocytes/macrophages) from total $CD45^+$ leukocytes for each peptide hydrogel at 3, 7, and 14 days-post injection. *Statistical comparison: Day 3, TLK5 vs TLE5 p-value 0.0044, TLE5 vs HLT2 and MAX8 p-value <0.0006. Day 7, MAX8 vs TLK5 and HLT2 p-value < 0.0296. Day 14, TLE5 vs TLK5 and MAX8 p-value < 0.0001, TLE5 vs HLT2 p-value 0.0002.* n = 5 mice. Error bars represent standard deviation.

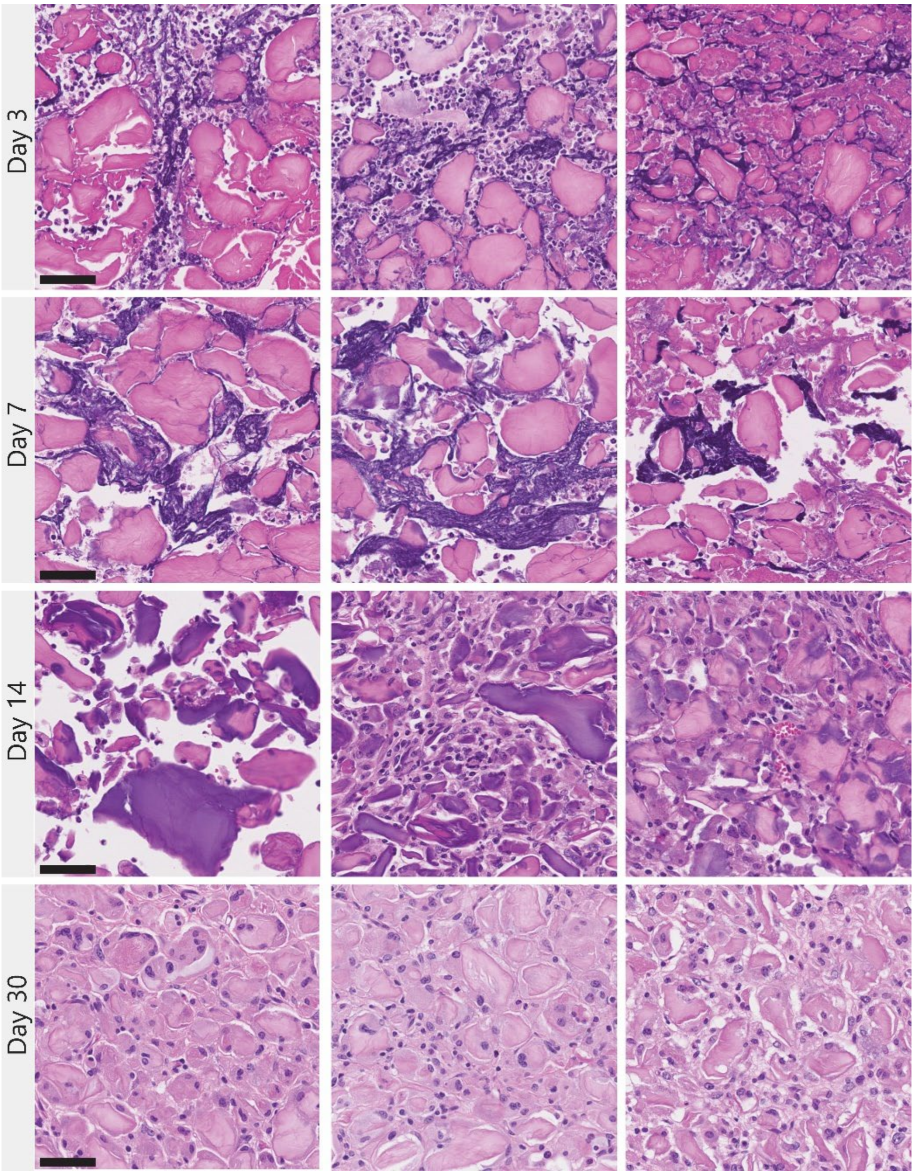

***Figure S14.*** H&E-stained tissue sections of TLK5 implants at different time points. At day 3 and 7 post-injections, the implants present areas with extracellular traps seen as basophilic DNA fibrous networks. Scale bar = 50 μm.

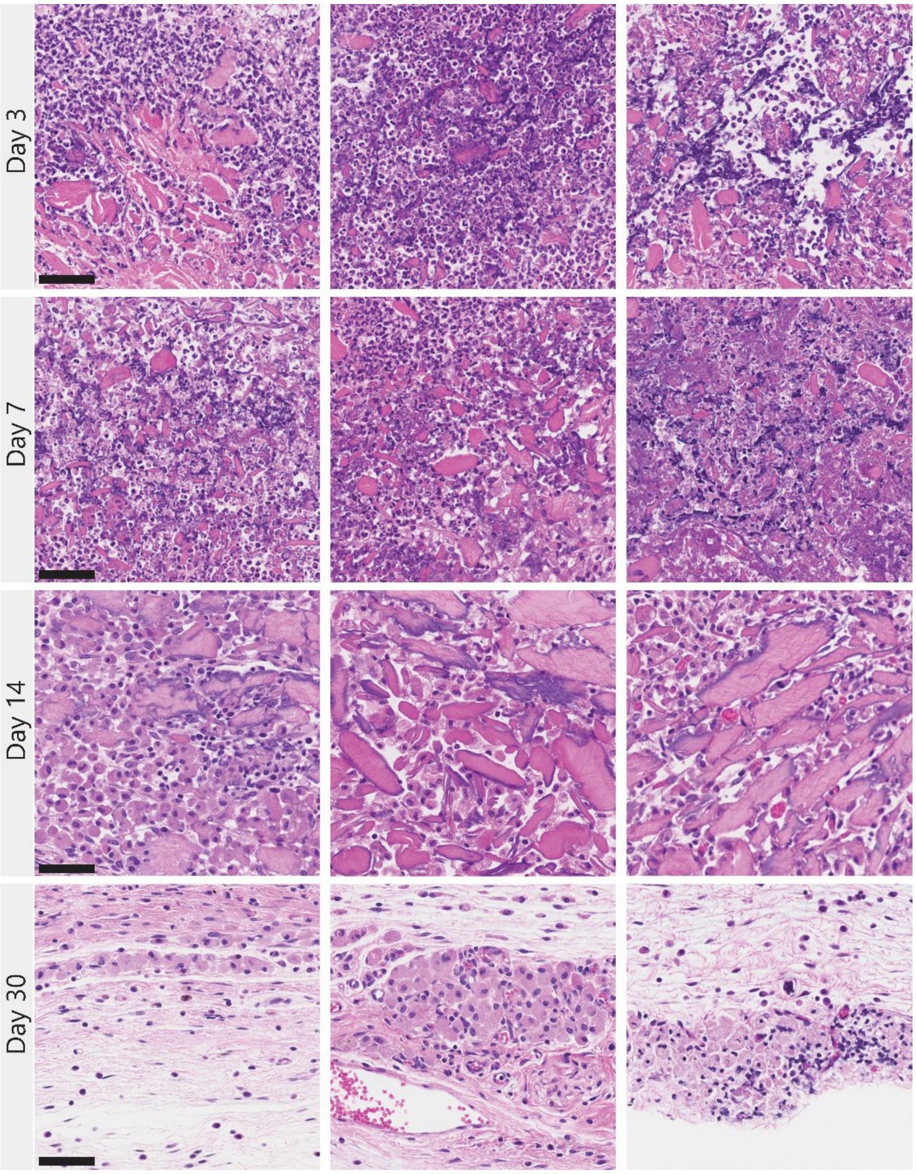

***Figure S15.*** H&E-stained tissue sections of HLT2 implants at different time points. At day 3 and 7 post-injections, the implants present areas with extracellular traps seen as basophilic DNA fibrous networks. Scale bar = 50 μm.

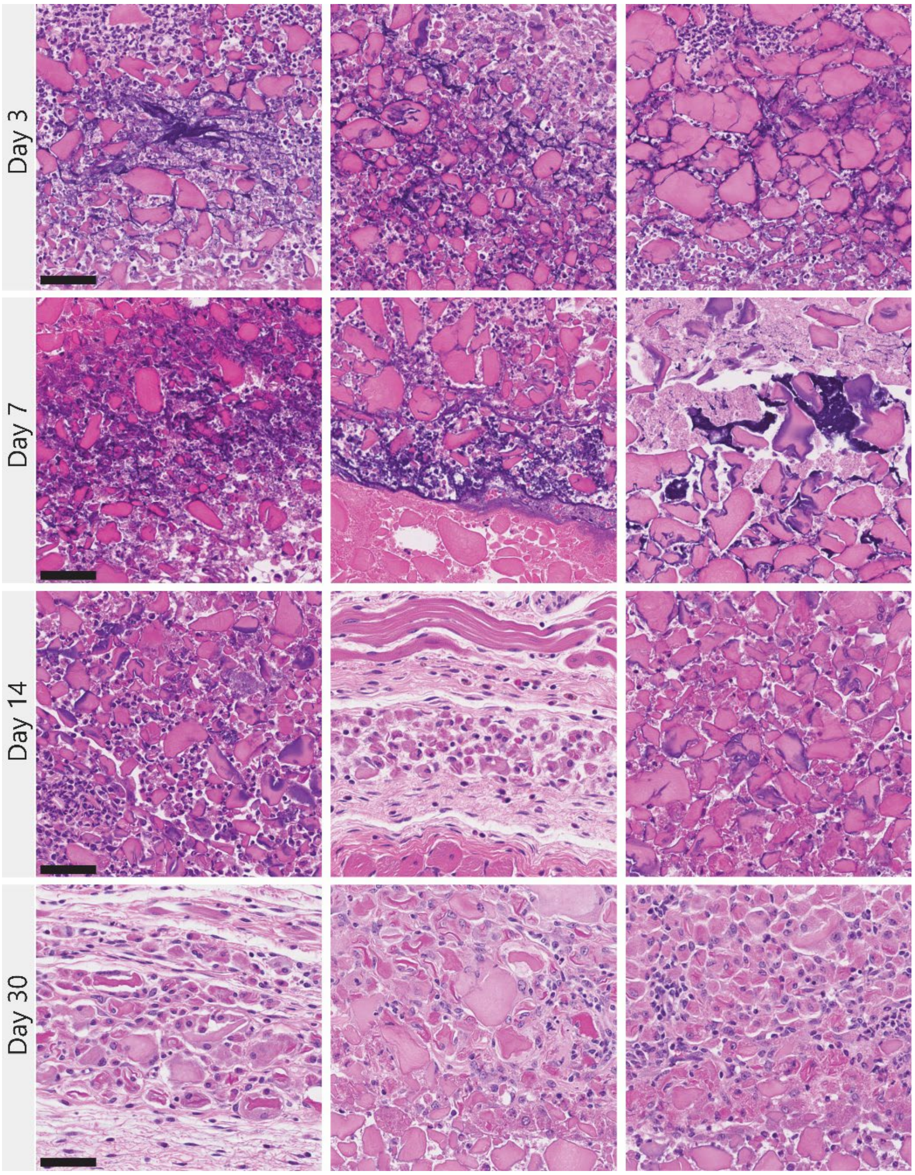

***Figure S16.*** H&E-stained tissue sections of MAX8 implants at different time points. At day 3 and 7 post-injections, the implants present areas with extracellular traps seen as basophilic DNA fibrous networks. Scale bar = 50 μm.

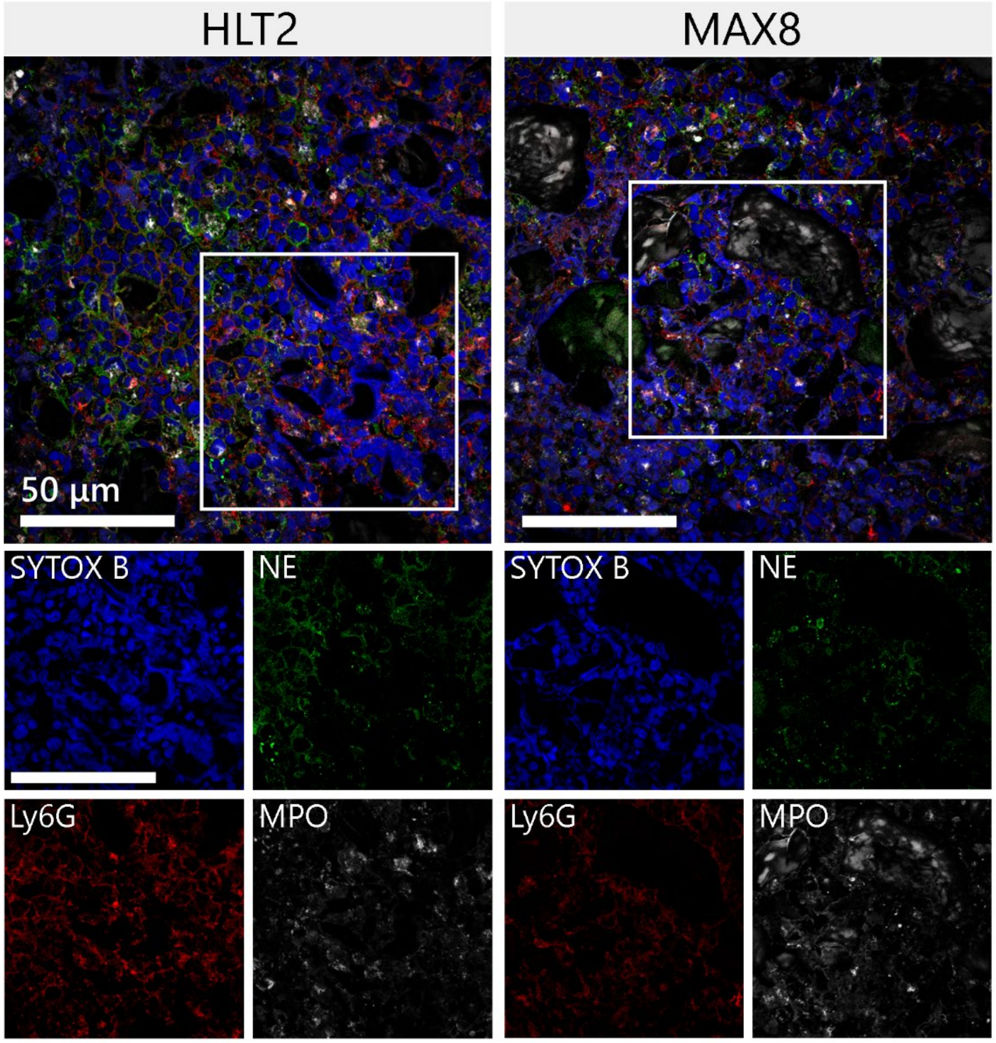

***Figure S17.*** Representative immunofluorescence images of HLT2 and MAX8 implants stained for NETs markers NE (green), MPO (gray), DNA (blue), and Ly6G (red) at day 3 post injection. Scale bar = 50 µm.

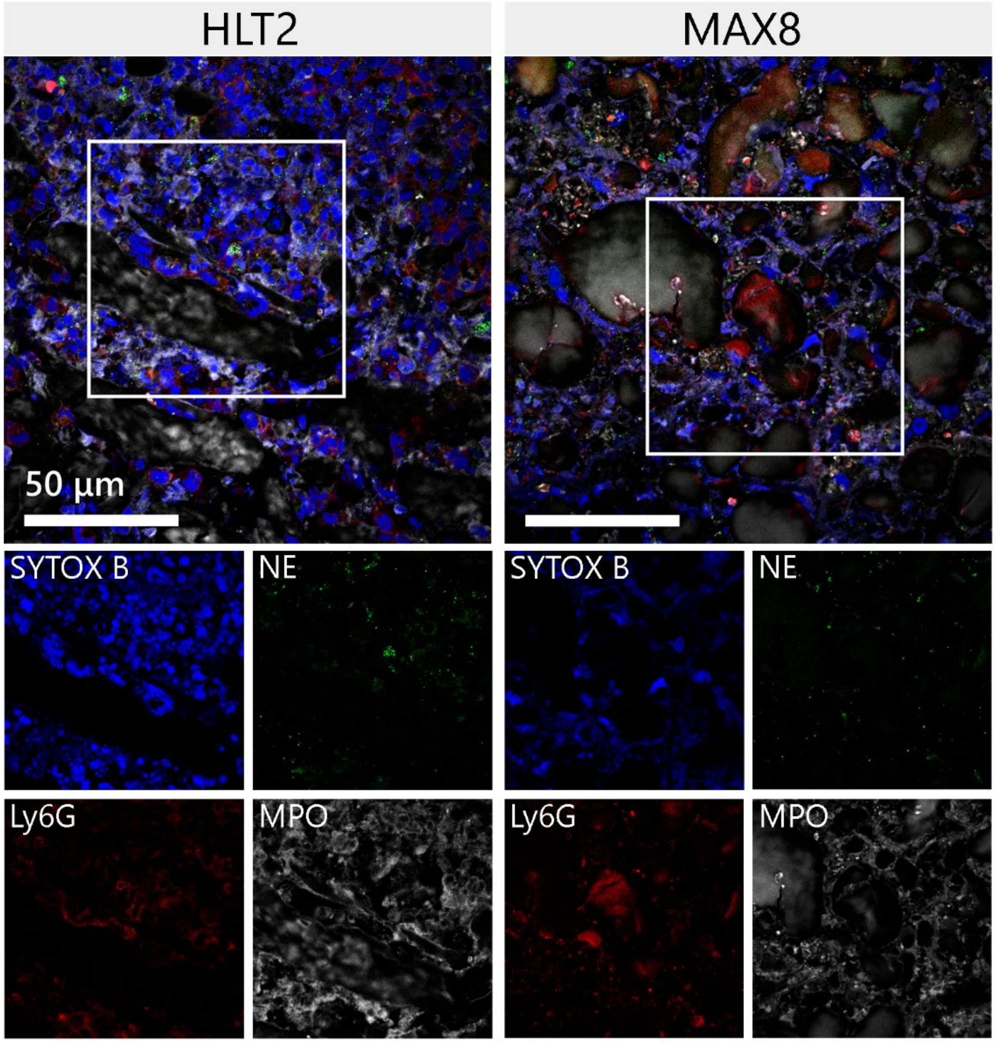

***Figure S18.*** Representative immunofluorescence images of HLT2 and MAX8 implants stained for NETs markers NE (green), MPO (gray), DNA (blue), and Ly6G (red) at day 7 post injection. Scale bar = 50 µm.

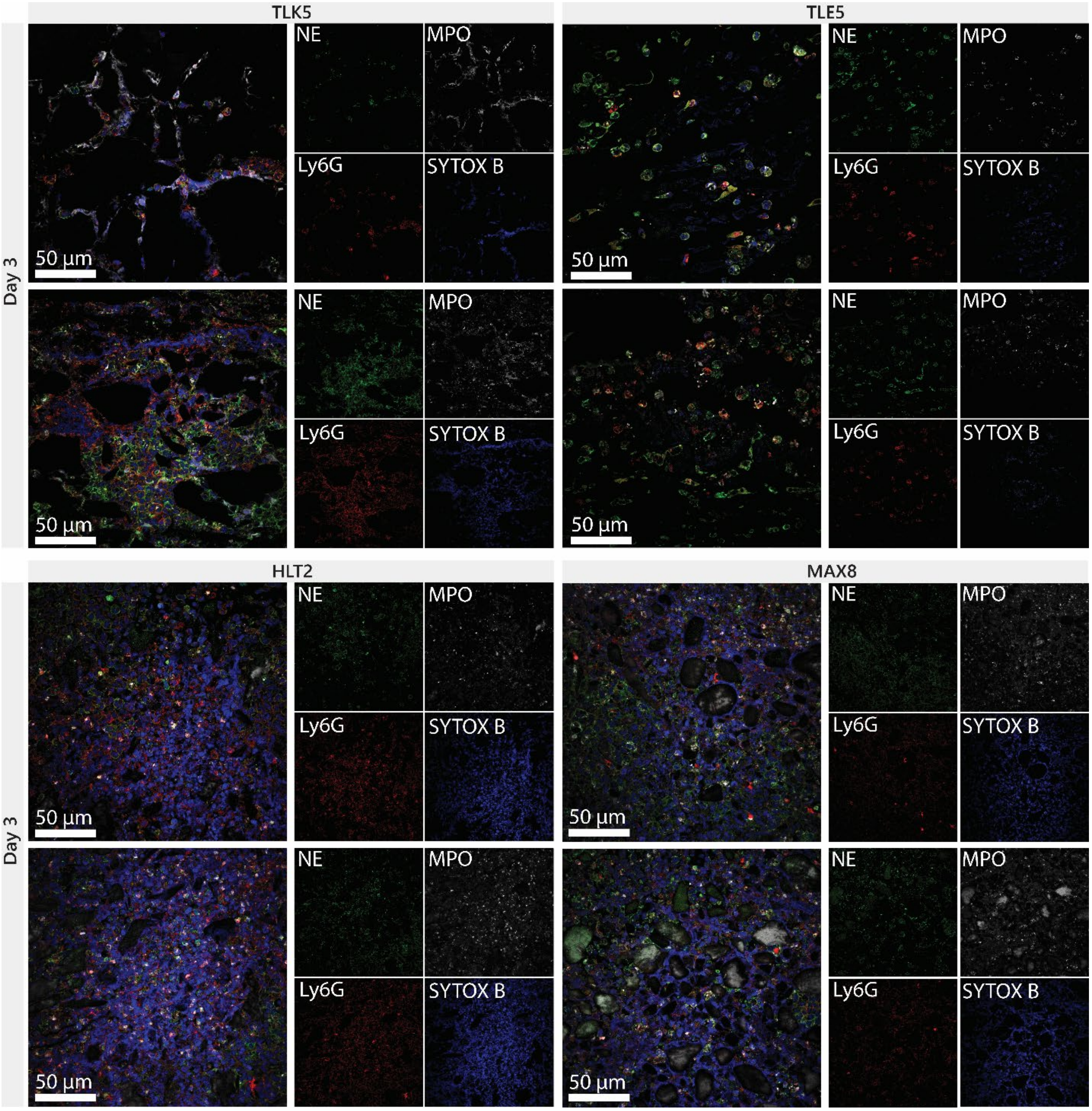

***Figure S19.*** Representative immunofluorescence images of different implants for TLK5, TLE5, HLT2, and MAX8 hydrogels three days post-injection.

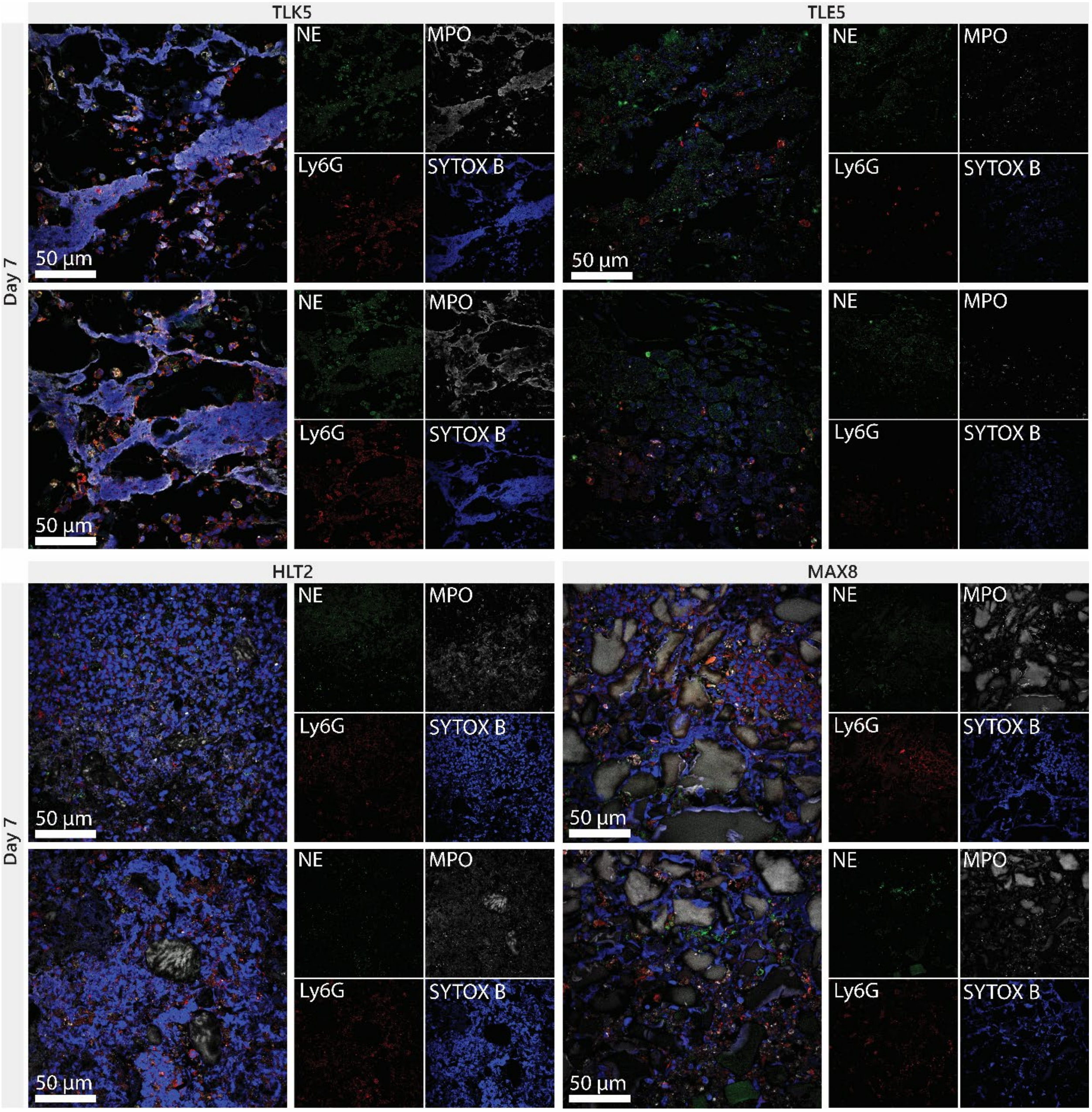

***Figure S20.*** Representative immunofluorescence images of different implants for TLK5, TLE5, HLT2, and MAX8 hydrogels seven days post-injection.

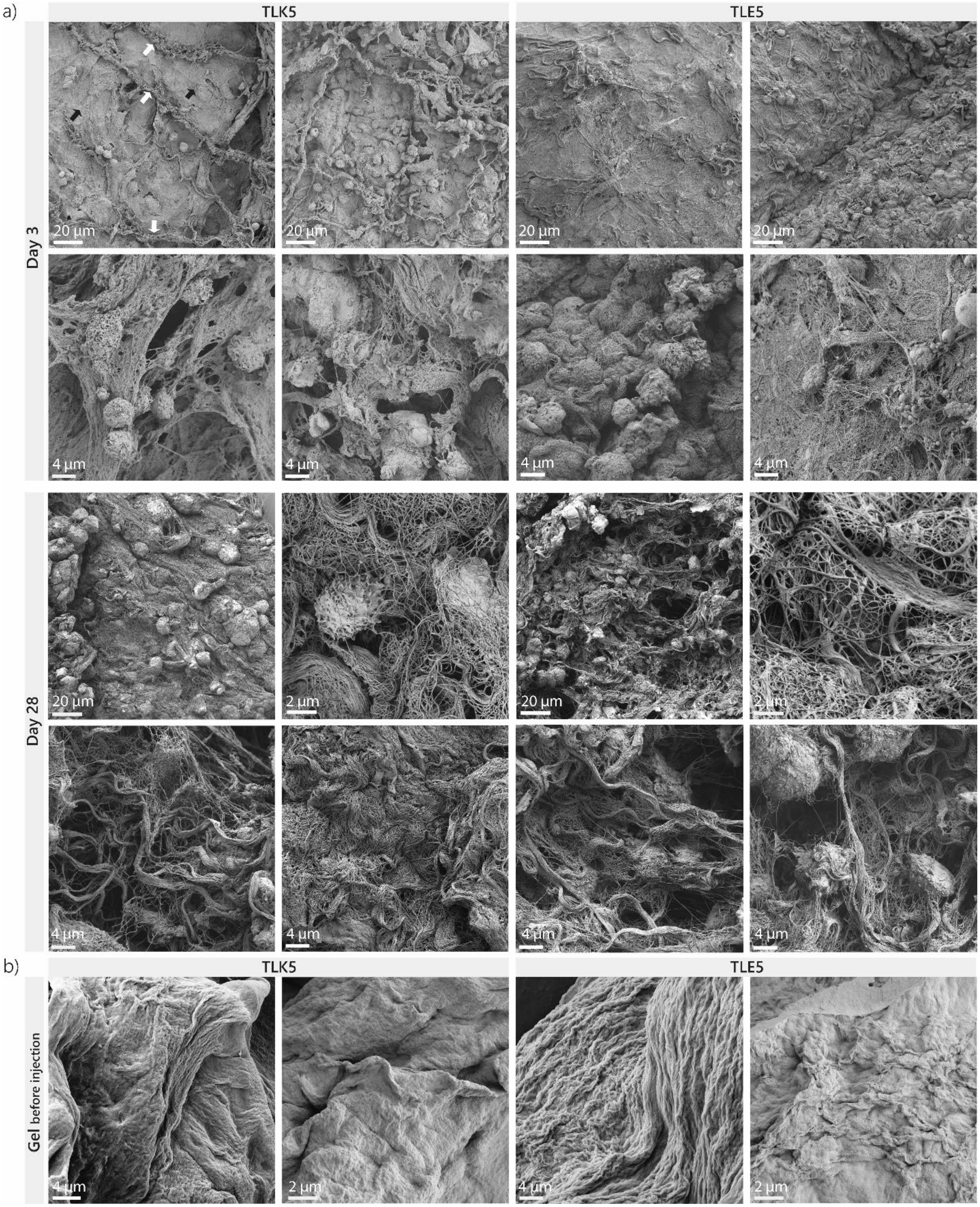

***Figure S21.*** SEM images of TLK5 and TLE5 gels. a) TLK5 and TLE5 implants injected in the subcutaneous space at day 3 and day 28 post-injection. TLK5 gel is shown as black arrows and NETs are shown as white arrows. b) SEM images of TLK5 and TLE5 native hydrogels before injection.

***Table S3.*** Statistical comparisons for Figure 5- Enzymes, cytokines, and chemokines.

*Neutrophil Elastase (NE) Day 1*

| Tukey's multiple comparisons test | Mean Diff. | 95.00% CI of diff. | Below threshold? | Summary | Adjusted P Value |
|---|---|---|---|---|---|
| TLK5 vs. HLT2 | -13883 | -33597 to 5832 | No | ns | 0.2236 |
| TLK5 vs. MAX8 | -11855 | -31570 to 7859 | No | ns | 0.3458 |
| TLK5 vs. TLE5 | 23674 | 3960 to 43389 | Yes | * | 0.0161 |
| HLT2 vs. MAX8 | 2027 | -17687 to 21742 | No | ns | 0.9908 |
| HLT2 vs. TLE5 | 37557 | 17843 to 57271 | Yes | *** | 0.0003 |
| MAX8 vs. TLE5 | 35530 | 15815 to 55244 | Yes | *** | 0.0005 |

*Neutrophil Elastase (NE) Day 3*

| Tukey's multiple comparisons test | Mean Diff. | 95.00% CI of diff. | Below threshold? | Summary | Adjusted P Value |
|---|---|---|---|---|---|
| TLK5 vs. HLT2 | -3603 | -20112 to 12906 | No | ns | 0.9227 |
| TLK5 vs. MAX8 | 4620 | -11889 to 21129 | No | ns | 0.8531 |
| TLK5 vs. TLE5 | 16015 | -494.5 to 32524 | No | ns | 0.0589 |
| HLT2 vs. MAX8 | 8223 | -8286 to 24732 | No | ns | 0.5027 |
| HLT2 vs. TLE5 | 19618 | 3109 to 36127 | Yes | * | 0.0173 |
| MAX8 vs. TLE5 | 11395 | -5114 to 27904 | No | ns | 0.2381 |

*Neutrophil Elastase (NE) Day 7*

| Tukey's multiple comparisons test | Mean Diff. | 95.00% CI of diff. | Below threshold? | Summary | Adjusted P Value |
|---|---|---|---|---|---|
| TLK5 vs. HLT2 | 3878 | -1442 to 9198 | No | ns | 0.1997 |
| TLK5 vs. MAX8 | 4928 | -392.3 to 10247 | No | ns | 0.0745 |
| TLK5 vs. TLE5 | 12887 | 7567 to 18207 | Yes | **** | <0.0001 |
| HLT2 vs. MAX8 | 1050 | -4270 to 6369 | No | ns | 0.9412 |
| HLT2 vs. TLE5 | 9009 | 3689 to 14329 | Yes | *** | 0.0009 |
| MAX8 vs. TLE5 | 7959 | 2639 to 13279 | Yes | ** | 0.0029 |

*Neutrophil Elastase (NE) Day 14*

| Tukey's multiple comparisons test | Mean Diff. | 95.00% CI of diff. | Below threshold? | Summary | Adjusted P Value |
|---|---|---|---|---|---|
| TLK5 vs. HLT2 | 877.2 | -2268 to 4022 | No | ns | 0.8543 |
| TLK5 vs. MAX8 | -6268 | -9413 to -3124 | Yes | *** | 0.0002 |
| TLK5 vs. TLE5 | -3293 | -6438 to -148.6 | Yes | * | 0.0385 |
| HLT2 vs. MAX8 | -7146 | -10290 to -4001 | Yes | **** | <0.0001 |
| HLT2 vs. TLE5 | -4171 | -7315 to -1026 | Yes | ** | 0.0078 |
| MAX8 vs. TLE5 | 2975 | -169.7 to 6120 | No | ns | 0.0670 |

*Myeloperoxidase (MPO) Day 1*

| Tukey's multiple comparisons test | Mean Diff. | 95.00% CI of diff. | Below threshold? | Summary | Adjusted P Value |
|---|---|---|---|---|---|
| TLK5 vs. HLT2 | 21765 | -24250 to 67780 | No | ns | 0.5445 |
| TLK5 vs. MAX8 | 43124 | -2891 to 89140 | No | ns | 0.0703 |
| TLK5 vs. TLE5 | 78306 | 32291 to 124321 | Yes | *** | 0.0009 |
| HLT2 vs. MAX8 | 21359 | -24656 to 67374 | No | ns | 0.5594 |
| HLT2 vs. TLE5 | 56541 | 10525 to 102556 | Yes | * | 0.0137 |
| MAX8 vs. TLE5 | 35181 | -10834 to 81197 | No | ns | 0.1690 |

*Myeloperoxidase (MPO) Day 3*

| Tukey's multiple comparisons test | Mean Diff. | 95.00% CI of diff. | Below threshold? | Summary | Adjusted P Value |
|---|---|---|---|---|---|
| TLK5 vs. HLT2 | 14210 | -19976 to 48395 | No | ns | 0.6420 |
| TLK5 vs. MAX8 | 8060 | -26125 to 42246 | No | ns | 0.9052 |
| TLK5 vs. TLE5 | 33653 | -532.5 to 67839 | No | ns | 0.0544 |
| HLT2 vs. MAX8 | -6149 | -40335 to 28037 | No | ns | 0.9544 |
| HLT2 vs. TLE5 | 19444 | -14742 to 53629 | No | ns | 0.3921 |
| MAX8 vs. TLE5 | 25593 | -8593 to 59779 | No | ns | 0.1822 |

*Myeloperoxidase (MPO) Day 7*

| Tukey's multiple comparisons test | Mean Diff. | 95.00% CI of diff. | Below threshold? | Summary | Adjusted P Value |
|---|---|---|---|---|---|
| TLK5 vs. HLT2 | 6673 | -48148 to 61495 | No | ns | 0.9850 |
| TLK5 vs. MAX8 | -10231 | -65053 to 44590 | No | ns | 0.9495 |
| TLK5 vs. TLE5 | 50556 | -4266 to 105377 | No | ns | 0.0761 |
| HLT2 vs. MAX8 | -16904 | -71726 to 37917 | No | ns | 0.8140 |
| HLT2 vs. TLE5 | 43883 | -10939 to 98704 | No | ns | 0.1420 |
| MAX8 vs. TLE5 | 60787 | 5965 to 115609 | Yes | * | 0.0272 |

*Myeloperoxidase (MPO) Day 14*

| Tukey's multiple comparisons test | Mean Diff. | 95.00% CI of diff. | Below threshold? | Summary | Adjusted P Value |
|---|---|---|---|---|---|
| TLK5 vs. HLT2 | -16442 | -83827 to 50942 | No | ns | 0.8963 |
| TLK5 vs. MAX8 | -107498 | -174883 to -40114 | Yes | ** | 0.0016 |
| TLK5 vs. TLE5 | -98723 | -166107 to -31338 | Yes | ** | 0.0035 |
| HLT2 vs. MAX8 | -91056 | -158441 to -23671 | Yes | ** | 0.0067 |
| HLT2 vs. TLE5 | -82280 | -149665 to -14896 | Yes | * | 0.0143 |
| MAX8 vs. TLE5 | 8776 | -58609 to 76160 | No | ns | 0.9817 |

*CXCL1 (KC) Day 1*

| Tukey's multiple comparisons test | Mean Diff. | 95.00% CI of diff. | Below threshold? | Summary | Adjusted P Value |
|---|---|---|---|---|---|
| TLK5 vs. HLT2 | -166.3 | -661.5 to 328.9 | No | ns | 0.7731 |
| TLK5 vs. MAX8 | -2.679 | -497.9 to 492.5 | No | ns | >0.9999 |
| TLK5 vs. TLE5 | 573.8 | 78.56 to 1069 | Yes | * | 0.0205 |
| HLT2 vs. MAX8 | 163.6 | -331.6 to 658.8 | No | ns | 0.7814 |
| HLT2 vs. TLE5 | 740.1 | 244.8 to 1235 | Yes | ** | 0.0029 |
| MAX8 vs. TLE5 | 576.5 | 81.24 to 1072 | Yes | * | 0.0199 |

*CXCL1 (KC) Day 3*

| Tukey's multiple comparisons test | Mean Diff. | 95.00% CI of diff. | Below threshold? | Summary | Adjusted P Value |
|---|---|---|---|---|---|
| TLK5 vs. HLT2 | 30.61 | -13.99 to 75.21 | No | ns | 0.2393 |
| TLK5 vs. MAX8 | 57.40 | 10.09 to 104.7 | Yes | * | 0.0153 |
| TLK5 vs. TLE5 | 60.11 | 15.51 to 104.7 | Yes | ** | 0.0071 |
| HLT2 vs. MAX8 | 26.79 | -20.51 to 74.10 | No | ns | 0.3913 |
| HLT2 vs. TLE5 | 29.51 | -15.10 to 74.11 | No | ns | 0.2666 |
| MAX8 vs. TLE5 | 2.714 | -44.59 to 50.02 | No | ns | 0.9983 |

*CXCL1 (KC) Day 7*

| Tukey's multiple comparisons test | Mean Diff. | 95.00% CI of diff. | Below threshold? | Summary | Adjusted P Value |
|---|---|---|---|---|---|
| TLK5 vs. HLT2 | 1.810 | -43.55 to 47.17 | No | ns | 0.9994 |
| TLK5 vs. MAX8 | -8.260 | -53.62 to 37.10 | No | ns | 0.9528 |
| TLK5 vs. TLE5 | -28.70 | -74.06 to 16.66 | No | ns | 0.3045 |
| HLT2 vs. MAX8 | -10.07 | -55.43 to 35.29 | No | ns | 0.9191 |
| HLT2 vs. TLE5 | -30.51 | -75.87 to 14.85 | No | ns | 0.2571 |
| MAX8 vs. TLE5 | -20.44 | -65.80 to 24.92 | No | ns | 0.5823 |

*CXCL1 (KC) Day 14*

| Tukey's multiple comparisons test | Mean Diff. | 95.00% CI of diff. | Below threshold? | Summary | Adjusted P Value |
|---|---|---|---|---|---|
| TLK5 vs. HLT2 | 3.369 | -216.5 to 223.2 | No | ns | TLK5 vs. HLT2 |
| TLK5 vs. MAX8 | -5.353 | -225.2 to 214.5 | No | ns | TLK5 vs. MAX8 |
| TLK5 vs. TLE5 | -326.8 | -546.6 to -106.9 | Yes | ** | TLK5 vs. TLE5 |
| HLT2 vs. MAX8 | -8.722 | -228.6 to 211.1 | No | ns | HLT2 vs. MAX8 |
| HLT2 vs. TLE5 | -330.1 | -550.0 to -110.3 | Yes | ** | HLT2 vs. TLE5 |
| MAX8 vs. TLE5 | -321.4 | -541.2 to -101.6 | Yes | ** | MAX8 vs. TLE5 |

*CCL3 (MIP-1α) Day 1*

| Tukey's multiple comparisons test | Mean Diff. | 95.00% CI of diff. | Below threshold? | Summary | Adjusted P Value |
|---|---|---|---|---|---|
| TLK5 vs. HLT2 | 217.4 | 57.22 to 377.6 | Yes | ** | 0.0065 |
| TLK5 vs. MAX8 | 85.00 | -75.18 to 245.2 | No | ns | 0.4502 |
| TLK5 vs. TLE5 | 277.7 | 117.5 to 437.8 | Yes | *** | 0.0007 |
| HLT2 vs. MAX8 | -132.4 | -292.6 to 27.78 | No | ns | 0.1247 |
| HLT2 vs. TLE5 | 60.26 | -99.92 to 220.4 | No | ns | 0.7083 |
| MAX8 vs. TLE5 | 192.7 | 32.48 to 352.8 | Yes | * | 0.0159 |

*CCL3 (MIP-1α) Day 3*

| Tukey's multiple comparisons test | Mean Diff. | 95.00% CI of diff. | Below threshold? | Summary | Adjusted P Value |
|---|---|---|---|---|---|
| TLK5 vs. HLT2 | 331.0 | -80.64 to 742.6 | No | ns | 0.1395 |
| TLK5 vs. MAX8 | 628.2 | 216.6 to 1040 | Yes | ** | 0.0024 |
| TLK5 vs. TLE5 | 836.1 | 424.5 to 1248 | Yes | *** | 0.0001 |
| HLT2 vs. MAX8 | 297.3 | -114.3 to 708.9 | No | ns | 0.2060 |
| HLT2 vs. TLE5 | 505.2 | 93.55 to 916.8 | Yes | * | 0.0138 |
| MAX8 vs. TLE5 | 207.9 | -203.7 to 619.5 | No | ns | 0.4913 |

*CCL3 (MIP-1α) Day 7*

| Tukey's multiple comparisons test | Mean Diff. | 95.00% CI of diff. | Below threshold? | Summary | Adjusted P Value |
|---|---|---|---|---|---|
| TLK5 vs. HLT2 | 311.5 | -64.05 to 687.0 | No | ns | 0.1229 |
| TLK5 vs. MAX8 | 678.5 | 303.0 to 1054 | Yes | *** | 0.0005 |
| TLK5 vs. TLE5 | 1431 | 1056 to 1807 | Yes | **** | <0.0001 |
| HLT2 vs. MAX8 | 367.0 | -8.519 to 742.6 | No | ns | 0.0566 |
| HLT2 vs. TLE5 | 1120 | 744.2 to 1495 | Yes | **** | <0.0001 |
| MAX8 vs. TLE5 | 752.7 | 377.2 to 1128 | Yes | *** | 0.0002 |

*CCL3 (MIP-1α) Day 14*

| Tukey's multiple comparisons test | Mean Diff. | 95.00% CI of diff. | Below threshold? | Summary | Adjusted P Value |
|---|---|---|---|---|---|
| TLK5 vs. HLT2 | 103.9 | -96.86 to 304.6 | No | ns | TLK5 vs. HLT2 |
| TLK5 vs. MAX8 | -153.3 | -354.0 to 47.46 | No | ns | TLK5 vs. MAX8 |
| TLK5 vs. TLE5 | -134.6 | -335.3 to 66.14 | No | ns | TLK5 vs. TLE5 |
| HLT2 vs. MAX8 | -257.1 | -457.8 to -56.40 | Yes | * | HLT2 vs. MAX8 |
| HLT2 vs. TLE5 | -238.4 | -439.1 to -37.71 | Yes | * | HLT2 vs. TLE5 |
| MAX8 vs. TLE5 | 18.69 | -182.0 to 219.4 | No | ns | MAX8 vs. TLE5 |

*CCL2 (MCP-1) Day 1*

| Tukey's multiple comparisons test | Mean Diff. | 95.00% CI of diff. | Below threshold? | Summary | Adjusted P Value |
|---|---|---|---|---|---|
| TLK5 vs. HLT2 | 41.59 | -73.06 to 156.2 | No | ns | 0.7304 |
| TLK5 vs. MAX8 | 72.06 | -42.59 to 186.7 | No | ns | 0.3099 |
| TLK5 vs. TLE5 | 203.3 | 88.65 to 317.9 | Yes | *** | 0.0006 |
| HLT2 vs. MAX8 | 30.47 | -84.17 to 145.1 | No | ns | 0.8709 |
| HLT2 vs. TLE5 | 161.7 | 47.06 to 276.3 | Yes | ** | 0.0048 |
| MAX8 vs. TLE5 | 131.2 | 16.59 to 245.9 | Yes | * | 0.0222 |

*CCL2 (MCP-1) Day 3*

| Tukey's multiple comparisons test | Mean Diff. | 95.00% CI of diff. | Below threshold? | Summary | Adjusted P Value |
|---|---|---|---|---|---|
| TLK5 vs. HLT2 | 2.286 | -54.44 to 59.01 | No | ns | 0.9994 |
| TLK5 vs. MAX8 | -8.003 | -64.72 to 48.72 | No | ns | 0.9770 |
| TLK5 vs. TLE5 | -233.4 | -290.2 to -176.7 | Yes | **** | <0.0001 |
| HLT2 vs. MAX8 | -10.29 | -67.01 to 46.43 | No | ns | 0.9533 |
| HLT2 vs. TLE5 | -235.7 | -292.4 to -179.0 | Yes | **** | <0.0001 |
| MAX8 vs. TLE5 | -225.4 | -282.2 to -168.7 | Yes | **** | <0.0001 |

*CCL2 (MCP-1) Day 7*

| Tukey's multiple comparisons test | Mean Diff. | 95.00% CI of diff. | Below threshold? | Summary | Adjusted P Value |
|---|---|---|---|---|---|
| TLK5 vs. HLT2 | 2.078 | -54.64 to 58.80 | No | ns | 0.9996 |
| TLK5 vs. MAX8 | -44.17 | -100.9 to 12.55 | No | ns | 0.1578 |
| TLK5 vs. TLE5 | -280.1 | -336.8 to -223.4 | Yes | **** | <0.0001 |
| HLT2 vs. MAX8 | -46.25 | -103.0 to 10.47 | No | ns | 0.1319 |
| HLT2 vs. TLE5 | -282.1 | -338.9 to -225.4 | Yes | **** | <0.0001 |
| MAX8 vs. TLE5 | -235.9 | -292.6 to -179.2 | Yes | **** | <0.0001 |

*CCL2 (MCP-1) Day 14*

| Tukey's multiple comparisons test | Mean Diff. | 95.00% CI of diff. | Below threshold? | Summary | Adjusted P Value |
|---|---|---|---|---|---|
| TLK5 vs. HLT2 | 25.30 | -42.87 to 93.47 | No | ns | 0.7168 |
| TLK5 vs. MAX8 | -74.13 | -142.3 to -5.956 | Yes | * | 0.0308 |
| TLK5 vs. TLE5 | -112.4 | -180.6 to -44.26 | Yes | ** | 0.0012 |
| HLT2 vs. MAX8 | -99.42 | -167.6 to -31.25 | Yes | ** | 0.0036 |
| HLT2 vs. TLE5 | -137.7 | -205.9 to -69.55 | Yes | *** | 0.0001 |
| MAX8 vs. TLE5 | -38.30 | -106.5 to 29.87 | No | ns | 0.4023 |

*IL-6 Day 1*

| Tukey's multiple comparisons test | Mean Diff. | 95.00% CI of diff. | Below threshold? | Summary | Adjusted P Value |
|---|---|---|---|---|---|
| TLK5 vs. HLT2 | 410.7 | -443.0 to 1264 | No | ns | 0.5309 |
| TLK5 vs. MAX8 | 991.3 | 137.6 to 1845 | Yes | * | 0.0202 |
| TLK5 vs. TLE5 | 2510 | 1656 to 3364 | Yes | **** | <0.0001 |
| HLT2 vs. MAX8 | 580.6 | -273.1 to 1434 | No | ns | 0.2489 |
| HLT2 vs. TLE5 | 2099 | 1246 to 2953 | Yes | **** | <0.0001 |
| MAX8 vs. TLE5 | 1519 | 664.9 to 2372 | Yes | *** | 0.0006 |

*IL-6 Day 3*

| Tukey's multiple comparisons test | Mean Diff. | 95.00% CI of diff. | Below threshold? | Summary | Adjusted P Value |
|---|---|---|---|---|---|
| TLK5 vs. HLT2 | 154.5 | 78.24 to 230.7 | Yes | *** | 0.0001 |
| TLK5 vs. MAX8 | 185.0 | 108.8 to 261.2 | Yes | **** | <0.0001 |
| TLK5 vs. TLE5 | 189.2 | 113.0 to 265.4 | Yes | **** | <0.0001 |
| HLT2 vs. MAX8 | 30.51 | -45.70 to 106.7 | No | ns | 0.6680 |
| HLT2 vs. TLE5 | 34.73 | -41.48 to 110.9 | No | ns | 0.5738 |
| MAX8 vs. TLE5 | 4.219 | -71.99 to 80.43 | No | ns | 0.9985 |

*IL-6 Day 7*

| Tukey's multiple comparisons test | Mean Diff. | 95.00% CI of diff. | Below threshold? | Summary | Adjusted P Value |
|---|---|---|---|---|---|
| TLK5 vs. HLT2 | 9.543 | -17.15 to 36.24 | No | ns | 0.7389 |
| TLK5 vs. MAX8 | -0.1570 | -26.85 to 26.54 | No | ns | >0.9999 |
| TLK5 vs. TLE5 | 8.797 | -17.90 to 35.49 | No | ns | 0.7826 |
| HLT2 vs. MAX8 | -9.700 | -36.40 to 17.00 | No | ns | 0.7294 |
| HLT2 vs. TLE5 | -0.7457 | -27.44 to 25.95 | No | ns | 0.9998 |
| MAX8 vs. TLE5 | 8.954 | -17.74 to 35.65 | No | ns | 0.7736 |

*IL-6 Day 14*

| Tukey's multiple comparisons test | Mean Diff. | 95.00% CI of diff. | Below threshold? | Summary | Adjusted P Value |
|---|---|---|---|---|---|
| TLK5 vs. HLT2 | -0.08951 | -3.882 to 3.703 | No | ns | 0.9999 |
| TLK5 vs. MAX8 | -0.1118 | -3.396 to 3.172 | No | ns | 0.9996 |
| TLK5 vs. TLE5 | -3.873 | -7.157 to -0.5887 | Yes | * | 0.0189 |
| HLT2 vs. MAX8 | -0.02232 | -3.815 to 3.770 | No | ns | >0.9999 |
| HLT2 vs. TLE5 | -3.783 | -7.576 to 0.008919 | No | ns | 0.0506 |
| MAX8 vs. TLE5 | -3.761 | -7.045 to -0.4768 | Yes | * | 0.0228 |

*TNF-α Day 1*

| Tukey's multiple comparisons test | Mean Diff. | 95.00% CI of diff. | Below threshold? | Summary | Adjusted P Value |
|---|---|---|---|---|---|
| TLK5 vs. HLT2 | 18.04 | -21.13 to 57.21 | No | ns | 0.5656 |
| TLK5 vs. MAX8 | 54.26 | 15.09 to 93.43 | Yes | ** | 0.0055 |
| TLK5 vs. TLE5 | 76.48 | 37.31 to 115.7 | Yes | *** | 0.0002 |
| HLT2 vs. MAX8 | 36.22 | -2.952 to 75.39 | No | ns | 0.0751 |
| HLT2 vs. TLE5 | 58.44 | 19.27 to 97.61 | Yes | ** | 0.0030 |
| MAX8 vs. TLE5 | 22.22 | -16.95 to 61.39 | No | ns | 0.3942 |

*TNF-α Day 3*

| Tukey's multiple comparisons test | Mean Diff. | 95.00% CI of diff. | Below threshold? | Summary | Adjusted P Value |
|---|---|---|---|---|---|
| TLK5 vs. HLT2 | 46.45 | 4.409 to 88.49 | Yes | * | 0.0279 |
| TLK5 vs. MAX8 | 62.38 | 20.34 to 104.4 | Yes | ** | 0.0031 |
| TLK5 vs. TLE5 | 174.4 | 132.4 to 216.4 | Yes | **** | <0.0001 |
| HLT2 vs. MAX8 | 15.93 | -26.11 to 57.97 | No | ns | 0.7039 |
| HLT2 vs. TLE5 | 128.0 | 85.92 to 170.0 | Yes | **** | <0.0001 |
| MAX8 vs. TLE5 | 112.0 | 69.99 to 154.1 | Yes | **** | <0.0001 |

*TNF-α Day 7*

| Tukey's multiple comparisons test | Mean Diff. | 95.00% CI of diff. | Below threshold? | Summary | Adjusted P Value |
|---|---|---|---|---|---|
| TLK5 vs. HLT2 | 6.283 | -29.61 to 42.17 | No | ns | 0.9577 |
| TLK5 vs. MAX8 | 19.49 | -16.40 to 55.37 | No | ns | 0.4310 |
| TLK5 vs. TLE5 | 106.5 | 70.58 to 142.4 | Yes | **** | <0.0001 |
| HLT2 vs. MAX8 | 13.20 | -22.69 to 49.09 | No | ns | 0.7220 |
| HLT2 vs. TLE5 | 100.2 | 64.29 to 136.1 | Yes | **** | <0.0001 |
| MAX8 vs. TLE5 | 86.98 | 51.09 to 122.9 | Yes | **** | <0.0001 |

*TNF-α Day 14*

| Tukey's multiple comparisons test | Mean Diff. | 95.00% CI of diff. | Below threshold? | Summary | Adjusted P Value |
|---|---|---|---|---|---|
| TLK5 vs. HLT2 | 2.125 | -5.840 to 10.09 | No | ns | 0.8697 |
| TLK5 vs. MAX8 | -9.195 | -17.16 to -1.230 | Yes | * | 0.0210 |
| TLK5 vs. TLE5 | -6.393 | -14.36 to 1.572 | No | ns | 0.1405 |
| HLT2 vs. MAX8 | -11.32 | -19.29 to -3.355 | Yes | ** | 0.0045 |
| HLT2 vs. TLE5 | -8.518 | -16.48 to -0.5527 | Yes | * | 0.0340 |
| MAX8 vs. TLE5 | 2.802 | -5.163 to 10.77 | No | ns | 0.7480 |

*IL-1β Day 1*

| Tukey's multiple comparisons test | Mean Diff. | 95.00% CI of diff. | Below threshold? | Summary | Adjusted P Value |
|---|---|---|---|---|---|
| TLK5 vs. HLT2 | -6.199 | -25.23 to 12.83 | No | ns | 0.7885 |
| TLK5 vs. MAX8 | -0.7954 | -19.83 to 18.24 | No | ns | 0.9994 |
| TLK5 vs. TLE5 | 6.683 | -12.35 to 25.72 | No | ns | 0.7491 |
| HLT2 vs. MAX8 | 5.403 | -13.63 to 24.44 | No | ns | 0.8478 |
| HLT2 vs. TLE5 | 12.88 | -6.152 to 31.91 | No | ns | 0.2526 |
| MAX8 vs. TLE5 | 7.478 | -11.56 to 26.51 | No | ns | 0.6805 |

*IL-1β Day 3*

| Tukey's multiple comparisons test | Mean Diff. | 95.00% CI of diff. | Below threshold? | Summary | Adjusted P Value |
|---|---|---|---|---|---|
| TLK5 vs. HLT2 | 20.17 | -0.1632 to 40.51 | No | ns | 0.0522 |
| TLK5 vs. MAX8 | 25.28 | 4.948 to 45.62 | Yes | * | 0.0126 |
| TLK5 vs. TLE5 | 48.79 | 28.46 to 69.13 | Yes | **** | <0.0001 |
| HLT2 vs. MAX8 | 5.112 | -15.22 to 25.45 | No | ns | 0.8880 |
| HLT2 vs. TLE5 | 28.62 | 8.285 to 48.95 | Yes | ** | 0.0049 |
| MAX8 vs. TLE5 | 23.51 | 3.173 to 43.84 | Yes | * | 0.0208 |

*IL-1β Day 7*

| Tukey's multiple comparisons test | Mean Diff. | 95.00% CI of diff. | Below threshold? | Summary | Adjusted P Value |
|---|---|---|---|---|---|
| TLK5 vs. HLT2 | 10.67 | 0.6229 to 20.71 | Yes | * | 0.0355 |
| TLK5 vs. MAX8 | 11.08 | 1.033 to 21.12 | Yes | * | 0.0282 |
| TLK5 vs. TLE5 | 31.60 | 21.55 to 41.64 | Yes | **** | <0.0001 |
| HLT2 vs. MAX8 | 0.4104 | -9.633 to 10.45 | No | ns | 0.9994 |
| HLT2 vs. TLE5 | 20.93 | 10.89 to 30.97 | Yes | *** | 0.0001 |
| MAX8 vs. TLE5 | 20.52 | 10.48 to 30.56 | Yes | *** | 0.0001 |

*IL-1β Day 14*

| Tukey's multiple comparisons test | Mean Diff. | 95.00% CI of diff. | Below threshold? | Summary | Adjusted P Value |
|---|---|---|---|---|---|
| TLK5 vs. HLT2 | -0.5344 | -2.685 to 1.616 | No | ns | 0.8891 |
| TLK5 vs. MAX8 | -0.6339 | -2.662 to 1.394 | No | ns | 0.8044 |
| TLK5 vs. TLE5 | -2.422 | -4.450 to -0.3947 | Yes | * | 0.0170 |
| HLT2 vs. MAX8 | -0.09952 | -2.250 to 2.051 | No | ns | 0.9991 |
| HLT2 vs. TLE5 | -1.888 | -4.038 to 0.2628 | No | ns | 0.0953 |
| MAX8 vs. TLE5 | -1.788 | -3.816 to 0.2393 | No | ns | 0.0933 |

*G-CSF Day 1*

| Tukey's multiple comparisons test | Mean Diff. | 95.00% CI of diff. | Below threshold? | Summary | Adjusted P Value |
|---|---|---|---|---|---|
| TLK5 vs. HLT2 | -1755 | -2764 to -746.8 | Yes | *** | 0.0007 |
| TLK5 vs. MAX8 | 1722 | 713.2 to 2730 | Yes | *** | 0.0009 |
| TLK5 vs. TLE5 | 2963 | 1954 to 3971 | Yes | **** | <0.0001 |
| HLT2 vs. MAX8 | 3477 | 2469 to 4486 | Yes | **** | <0.0001 |
| HLT2 vs. TLE5 | 4718 | 3709 to 5727 | Yes | **** | <0.0001 |
| MAX8 vs. TLE5 | 1241 | 232.1 to 2249 | Yes | * | 0.0136 |

*G-CSF Day 3*

| Tukey's multiple comparisons test | Mean Diff. | 95.00% CI of diff. | Below threshold? | Summary | Adjusted P Value |
|---|---|---|---|---|---|
| TLK5 vs. HLT2 | 283.6 | -314.9 to 882.1 | No | ns | 0.5432 |
| TLK5 vs. MAX8 | 512.8 | -85.66 to 1111 | No | ns | 0.1070 |
| TLK5 vs. TLE5 | 797.8 | 199.3 to 1396 | Yes | ** | 0.0075 |
| HLT2 vs. MAX8 | 229.3 | -369.2 to 827.8 | No | ns | 0.6969 |
| HLT2 vs. TLE5 | 514.2 | -84.26 to 1113 | No | ns | 0.1057 |
| MAX8 vs. TLE5 | 285.0 | -313.5 to 883.5 | No | ns | 0.5393 |

*G-CSF Day 7*

| Tukey's multiple comparisons test | Mean Diff. | 95.00% CI of diff. | Below threshold? | Summary | Adjusted P Value |
|---|---|---|---|---|---|
| TLK5 vs. HLT2 | -23.87 | -123.3 to 75.52 | No | ns | 0.8985 |
| TLK5 vs. MAX8 | -23.14 | -122.5 to 76.26 | No | ns | 0.9064 |
| TLK5 vs. TLE5 | 29.04 | -70.35 to 128.4 | No | ns | 0.8337 |
| HLT2 vs. MAX8 | 0.7327 | -92.98 to 94.45 | No | ns | >0.9999 |
| HLT2 vs. TLE5 | 52.92 | -40.79 to 146.6 | No | ns | 0.3937 |
| MAX8 vs. TLE5 | 52.19 | -41.53 to 145.9 | No | ns | 0.4053 |

*G-CSF Day 14*

| Tukey's multiple comparisons test | Mean Diff. | 95.00% CI of diff. | Below threshold? | Summary | Adjusted P Value |
|---|---|---|---|---|---|
| TLK5 vs. HLT2 | -4.705 | -690.7 to 681.3 | No | ns | >0.9999 |
| TLK5 vs. MAX8 | 1.566 | -684.5 to 687.6 | No | ns | >0.9999 |
| TLK5 vs. TLE5 | -956.1 | -1642 to -270.0 | Yes | ** | 0.0053 |
| HLT2 vs. MAX8 | 6.271 | -679.7 to 692.3 | No | ns | >0.9999 |
| HLT2 vs. TLE5 | -951.4 | -1637 to -265.3 | Yes | ** | 0.0055 |
| MAX8 vs. TLE5 | -957.6 | -1644 to -271.6 | Yes | ** | 0.0052 |

*CCL5 RANTES Day 1*

| Tukey's multiple comparisons test | Mean Diff. | 95.00% CI of diff. | Below threshold? | Summary | Adjusted P Value |
|---|---|---|---|---|---|
| TLK5 vs. HLT2 | 0.5697 | -1.973 to 3.112 | No | ns | 0.9171 |
| TLK5 vs. MAX8 | -4.562 | -7.104 to -2.019 | Yes | *** | 0.0005 |
| TLK5 vs. TLE5 | -1.219 | -3.762 to 1.323 | No | ns | 0.5335 |
| HLT2 vs. MAX8 | -5.131 | -7.674 to -2.589 | Yes | *** | 0.0002 |
| HLT2 vs. TLE5 | -1.789 | -4.332 to 0.7536 | No | ns | 0.2242 |
| MAX8 vs. TLE5 | 3.342 | 0.7994 to 5.885 | Yes | ** | 0.0083 |

*CCL5 RANTES Day 3*

| Tukey's multiple comparisons test | Mean Diff. | 95.00% CI of diff. | Below threshold? | Summary | Adjusted P Value |
|---|---|---|---|---|---|
| TLK5 vs. HLT2 | 0.01392 | -0.5559 to 0.5837 | No | ns | 0.9999 |
| TLK5 vs. MAX8 | -0.5455 | -1.115 to 0.02428 | No | ns | 0.0630 |
| TLK5 vs. TLE5 | -0.2948 | -0.8646 to 0.2750 | No | ns | 0.4713 |
| HLT2 vs. MAX8 | -0.5594 | -1.129 to 0.01035 | No | ns | 0.0552 |
| HLT2 vs. TLE5 | -0.3087 | -0.8785 to 0.2611 | No | ns | 0.4327 |
| MAX8 vs. TLE5 | 0.2507 | -0.3191 to 0.8205 | No | ns | 0.6005 |

*CCL5 RANTES Day 7*

| Tukey's multiple comparisons test | Mean Diff. | 95.00% CI of diff. | Below threshold? | Summary | Adjusted P Value |
|---|---|---|---|---|---|
| TLK5 vs. HLT2 | 0.09916 | -1.737 to 1.935 | No | ns | 0.9986 |
| TLK5 vs. MAX8 | -3.497 | -5.333 to -1.662 | Yes | *** | 0.0003 |
| TLK5 vs. TLE5 | 0.2869 | -1.549 to 2.123 | No | ns | 0.9692 |
| HLT2 vs. MAX8 | -3.596 | -5.432 to -1.761 | Yes | *** | 0.0002 |
| HLT2 vs. TLE5 | 0.1878 | -1.648 to 2.023 | No | ns | 0.9909 |
| MAX8 vs. TLE5 | 3.784 | 1.948 to 5.620 | Yes | *** | 0.0001 |

*CCL5 RANTES Day 14*

| Tukey's multiple comparisons test | Mean Diff. | 95.00% CI of diff. | Below threshold? | Summary | Adjusted P Value |
|---|---|---|---|---|---|
| TLK5 vs. HLT2 | -7.483 | -15.36 to 0.3952 | No | ns | 0.0657 |
| TLK5 vs. MAX8 | -10.56 | -18.44 to -2.682 | Yes | ** | 0.0072 |
| TLK5 vs. TLE5 | 4.121 | -3.757 to 12.00 | No | ns | 0.4622 |
| HLT2 vs. MAX8 | -3.077 | -10.95 to 4.801 | No | ns | 0.6844 |
| HLT2 vs. TLE5 | 11.60 | 3.726 to 19.48 | Yes | ** | 0.0033 |
| MAX8 vs. TLE5 | 14.68 | 6.803 to 22.56 | Yes | *** | 0.0004 |

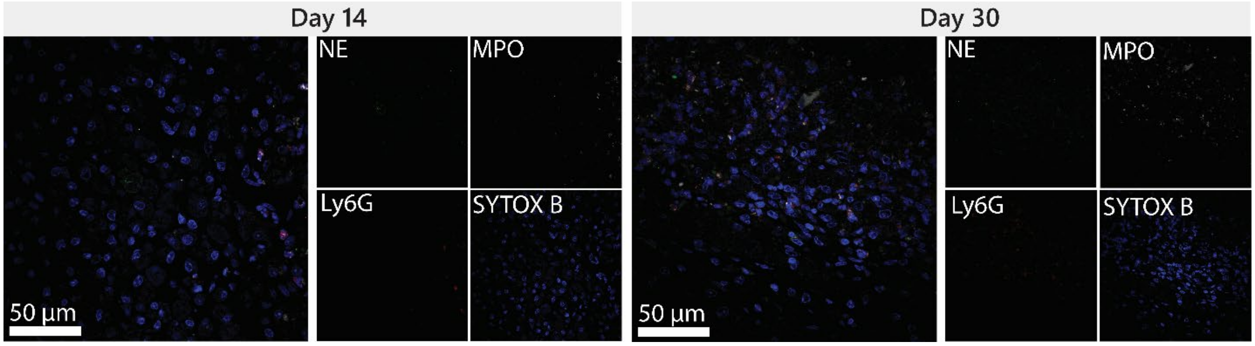

***Figure S22.*** Immunofluorescence staining of TLE5 implants 14- and 30-days post-injection.

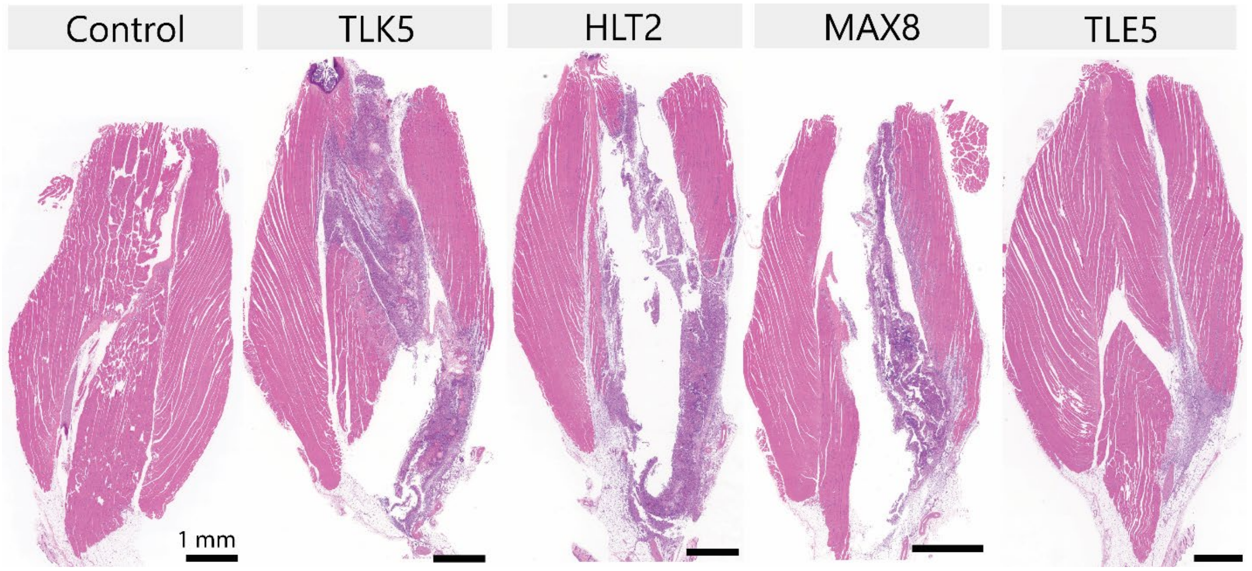

***Figure S23.*** H&E-stained tissue sections of the gastrocnemius muscle with gel injections.

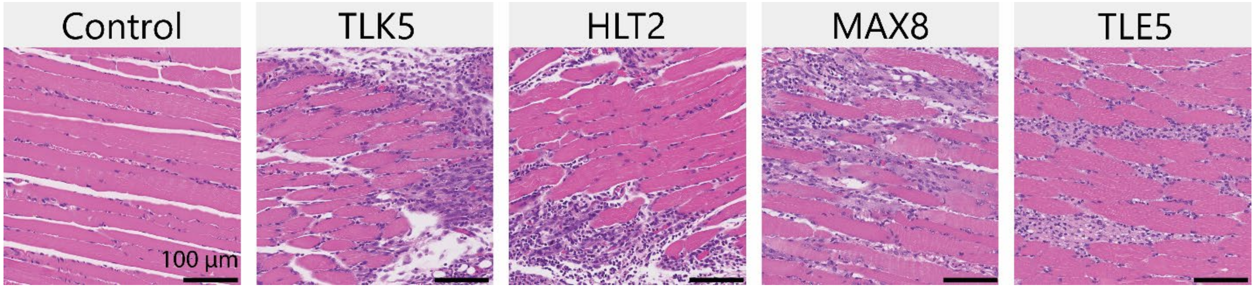

***Figure S24.*** H&E-stained tissue sections of muscle near the gel implant.
